# Conserved Link between Catalytic Site Interactions and Global Conformation in P-loop Enzymes

**DOI:** 10.1101/2022.07.13.499785

**Authors:** Fatlum Hajredini, Ranajeet Ghose

## Abstract

P-loop enzymes, ubiquitous in all of life’s domains and viruses, comprise a monophyletic group with pre-LUCA origins that have differentiated into several three-layered *α*/*β*/*α−* sandwich domain families utilizing a basic *β−* loop*−α−β* structural module housing conserved nucleotide-binding Walker-A and Walker-B sequences. We have analyzed a large dataset of P-loop enzyme structures representing both their KG and ASCE branches as proxies for their sampled conformational landscapes. We developed a novel framework to correlate global conformations and local catalytic site geometry, specifically involving the Walker motifs, to identify conserved signatures despite substantial structural and functional diversity. Our results suggest that P-loop enzymes populate global states broadly classifiable as open or closed. In the closed states, that share similar overall geometries irrespective of family, key catalytic site residues are aligned to optimally engage the critical Mg^2+^ ion suggesting compatibility with the chemical step. These catalytic site interactions are disrupted in the open states resulting in the loss of the Mg^2+^- coordinating ability yielding conformations incapable of chemistry. In contrast to the closed states, open states are highly diverse, and this variability is facilitated by differential coupling of specific residues that are part of, or spatially proximal to, the Walker motifs with the clade-specific tertiary fold. We suggest that an essential feature in the activation and nucleotide exchange processes for all P-loop enzymes is the universal coupling between global closure and local reorganization of the catalytic site for efficient coordination of Mg^2+^ that carries a tightly associated cargo, the substrate NTP.

## Introduction

The ancient phosphate-binding loop (P-loop) fold (1) that emerged before the last universal common ancestor (LUCA) is pervasive in all domains of life (2–4) and is also found in many viruses (5). P-loop enzymes (6) encode specific sequence signatures known as the Walker-A (GXXXXGK[T/S], where X is any residue) and Walker-B (*ϕϕϕϕ*D, where *ϕ* is a hydrophobic residue) motifs (7) organized within a minimal *β−*loop*−α−β* structural unit (Fig. 1A) that forms part of a 3-layered *α−β−α* sandwich. Within this unit, a glycine that initiates the Walker-A motif (Gly_A_) lies at the tip of a *β*-strand (*β*1) and the terminal lysine (Lys_A_) and serine/threonine (Ser_A_/Thr_A_) residues comprise the first two positions of the *α*-helix (*α*^P^). The Walker-B motif containing the aspartate (Asp_B_) lies on a separate *β*-strand (*β*3) that is spatially adjacent to *α*^P^. P-loop enzymes have enhanced this minimal module by acquiring a variety of structural modifications and decorations while refining their nucleoside triphosphate (NTP) specificity and tuning their ability to transfer the corresponding *γ*-phosphate to a variety of substrates including water (hydrolysis; NTPases) (3) and small molecules (phosphorylation; kinases) e.g., sugars, nucleotides etc. (4). In contrast, phosphorylation of tyrosine residues on protein substrates is generally considered to be exclusive to eukaryotes (8, 9), and catalyzed by protein tyrosine kinases with sequence (10) and structural signatures (11) distinct from those of the P-loop enzymes. This functional orthogonality defined by the eukaryotic protein tyrosine kinase (ePTK) and P-loop folds is broken by a family of enzymes, the BY-kinases, that are universally conserved in the bacterial kingdom (12, 13). The BY-kinase catalytic domain (CD) deploys a P-loop fold that encodes variations on the Walker-A and Walker-B motifs to facilitate tyrosine phosphorylation suggesting that this ability evolved in a fashion orthogonal to that in eukaryotes (9).

**Fig. 1.**
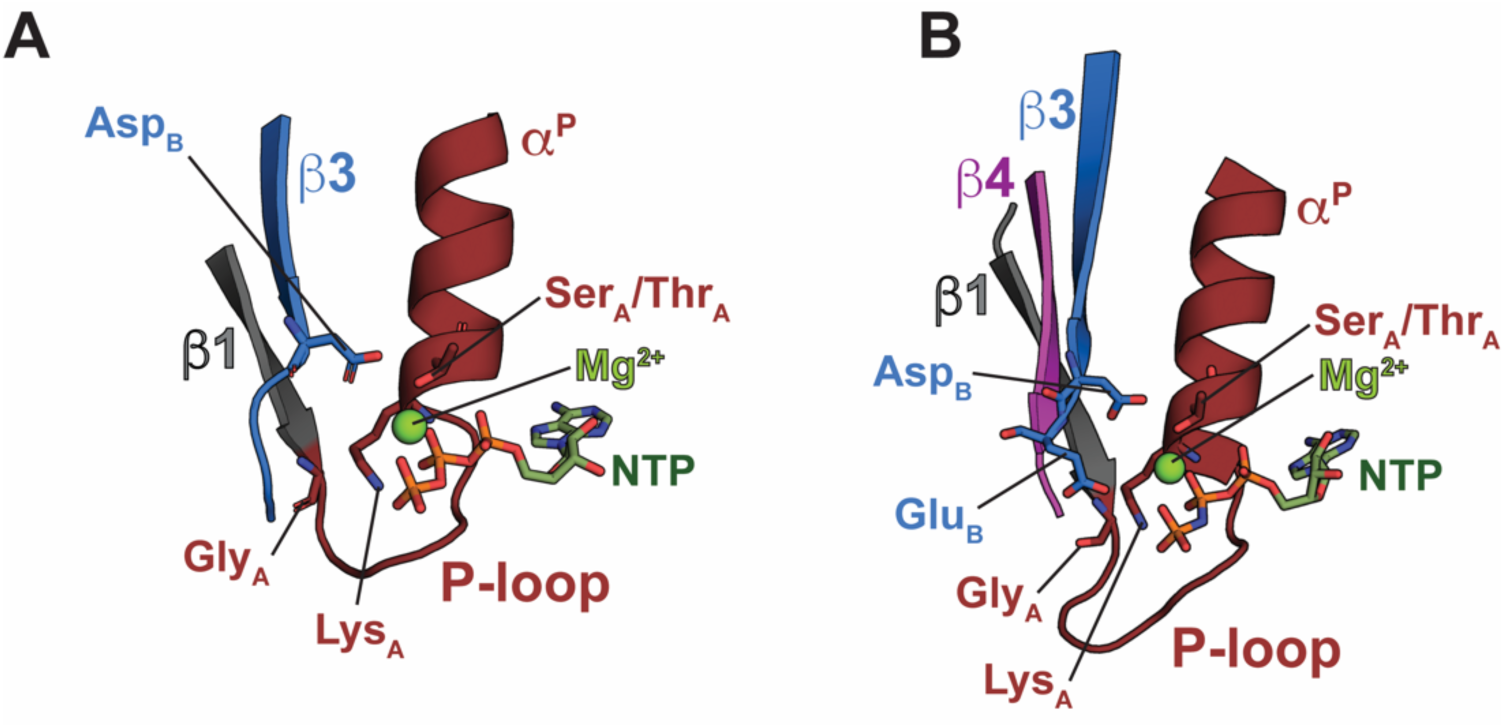
Minimal catalytic site architecture of P-loop enzymes. **(A)** A minimal *β*1*−*P-loop*−α*^P^*−β*3 unit is representative of the <u>k</u>inase <u>G</u>TPase (KG) branch of P-loop enzymes. The first Gly (Gly_A_) of the Walker-A motif that lies at the tip of a *β*-strand (*β*1), the phosphate-loop (P-loop), the *α*-helix (*α*^P^) bearing the catalytic Lys (Lys_A_) and the terminal Ser/Thr (Ser_A_/Thr_A_) of the Walker-A motif are indicated. Also shown is the Walker-B Asp (Asp_B_) that lies on a spatially proximal *β*-strand (*β*3). **(B)** The <u>a</u>dditional <u>s</u>trand <u>c</u>atalytic <u>E</u> (ASCE) branch contains an intervening strand (*β*4) sandwiched between *β*1 and *β*3, in addition to a “catalytic” Glu immediately following Asp_B_.

To uncover structural/conformational features that enable the somewhat unconventional use of the P-loop fold, we had previously investigated the mechanisms utilized by the CD of the *E. coli* BY-kinase Wzc to phosphorylate on tyrosine. We discovered a unique coupling between local interactions at the active site with structural transitions between “open” and “closed” states (*SI Appendix* Fig. S1A) that serves to temporally regulate a sequence of events that includes oligomerization, ATP binding, exclusion of bulk solvent, and *in trans* autophosphorylation (14, 15). The closed state was found to be characterized by an interaction between the conserved Walker residues, Thr_A_ and Asp_B_ (*SI Appendix* Fig. S1A), resulting in a configuration necessary to co-ordinate the catalytic Mg^2+^ (6). In contrast, in the open state, Asp_B_ was found to interact with the “catalytic” Lys_A_ yielding an inactive conformation incapable of Mg^2+^ coordination. Indeed, Mg^2+^ was able to drive transitions between the open and closed states by reconfiguring these interactions involving key active site elements (14, 15). We found similar Mg^2+^-induced, Walker-A/Walker-B-centered transitions between open and closed states in our computational studies on the P-loop containing shikimate kinase (SK) family (16). Indeed, survey of the literature suggests similar features in other P-loop enzymes e.g., the P-loop ATPase, Get3, also appears to populate compact (closed) and extended (open) structures in a manner correlated with active-site interactions (*SI Appendix* Fig. S1B) in the presence or absence of Mg^2+^ (and nucleotide) (17–19). Given these observations, we wondered whether these correlations between open/closed overall conformations, reshuffling of the interactions at the active site, and the presence/absence of the essential Mg^2+^ represent a fundamental feature in the conformational landscape of the P-loop enzymes. Addressing this issue requires a systematic analysis of a broad class of P-loop enzymes within a self-consistent framework, the goal of the work described here.

P-loop enzymes may be broadly classified into two major branches of related, but structurally and functionally diverse, proteins (4). The first, that includes small molecule kinases, GTPases and related ATPases, comprise the <u>k</u>inase <u>G</u>TPase (KG) (4) branch, of which Wzc, the SKs and Get3, mentioned above, are all members. The second, comprises the so-called <u>a</u>dditional <u>s</u>trand <u>c</u>atalytic Glu (ASCE) ATPases, that encode an additional *β*-strand sandwiched between *β*1 and the Walker-B bearing *β*3 (Fig. 1B). To test the universality of the structural coupling, described above, we developed a formalism that was applied to a large dataset comprising medium-to-high resolution structures of P-loop enzymes from both the KG and ASCE branches. Our analyses reveal that the open and closed states characterized by the relative orientation of key catalytic site elements appears to be a general feature of the P-loop enzymes. The open conformations are almost exclusively Mg^2+^-free, while the closed conformations are more likely to be Mg^2+^-bound. Further, while the closed states are similar in overall geometry, the open states are highly diverse and related to the specific structural fold and the resultant functional context. Based on our observations, we propose that, as demonstrated experimentally for Wzc (14), these open and closed states differ in their inherent abilities to optimally engage Mg^2+^, an essential co-factor of the NTP substrate, and therefore represent inactive and chemistry-compatible states, respectively. We further suggest these Mg^2+^-driven transitions constitutes a general mechanism of nucleotide exchange that is fine-tuned to the specific functional context of a given P-loop enzyme.

## Results

### Definition of a generalized coordinate system to characterize short- and long-range structural coupling in P-loop enzymes

As described above, experimental, and computational studies suggest the presence of open and closed conformations in specific P-loop enzymes. To establish whether these represent a more general characteristic, a set of parameters (generalized coordinates) to define these states is needed. A generalized coordinate system that relies on structural features that are conserved in all P-loop enzymes, rather than on class/family-specific characteristics, can be defined as shown in Fig. 2 (see Materials and Methods for additional details). This reference frame, that mirrors conventional spherical polar coordinates, is represented by a radial coordinate, *D*^W^, defined as Euclidian distance between the Ser_A_/Thr_A_ C*β* and Asp_B_ C*γ*, and angular coordinates, *ϕ* and *θ*, that specify the orientation of the Walker-B motif relative to *α*^P^. Given that the *D*^W^*−θ−ϕ* space relates sidechain conformations of key active site residues to the overall geometry of the core, the latter reflecting differences in the alignment of key segments of secondary structure between open and closed states, we refer to these states as “global” conformations.

**Fig. 2.**
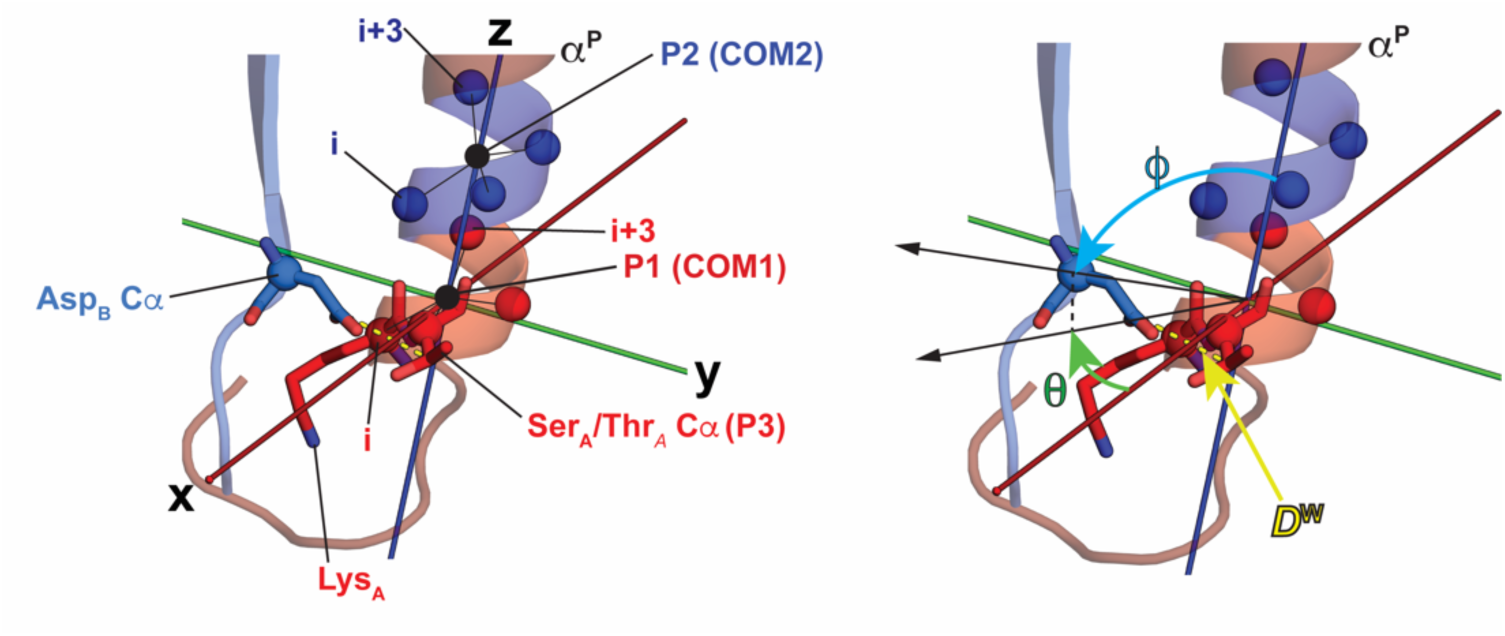
Definition of the spherical coordinate frame. The points used to define the spherical coordinate frame are shown on the left panel. The origin is defined by the center-of-mass (COM1, P1) of the C*α* atoms (red spheres) of residues that comprise the first turn of *α*^P^ starting with Lys_A_ (i, in red) and ending with i+3. The center-of-mass (COM2) of the C*α* atoms (dark-blue spheres) of residues of the second turn of *α*^P^ (i to i+3 in dark-blue) is used to define P2. P3 represents the C*α* atom of Ser_A_/Thr_A_. The definitions of *D*^W^, *ϕ* and *θ* are shown on the right panel. Cyan and green arrows indicate the directions of increase for the *ϕ* and *θ* variables, respectively. *D*^W^ (yellow dashes) measures the distance between the C*γ* of Asp_B_ (or C*δ* if a Glu is present at this position) and the C*β* atom of Ser_A_/Thr_A_.

We had previously used a two-dimensional cylindrical coordinate system (*SI Appendix* Fig. S2) comprising of a rise, |*h*|, and an angle, *θ* (here referred to as *θ*_cyl_ for clarity), to characterize the conformational ensembles of the CD of the BY-kinase, Wzc, obtained from molecular dynamics (MD) simulations (14). However, that reference frame is unsuitable for a general P-loop enzyme since its definition relies on structural elements specific to the BY-kinase family (*SI Appendix* Fig. S2). Given that the cylindrical coordinate system successfully captured the open and closed states of the Wzc CD (14, 16), an appropriate validation of the newly defined reference frame would comprise a demonstration of its ability to also parse out these two distinct global conformations. We projected the MD-generated ensemble of the ATP•Mg^2+^ complex of catalytic core of Wzc onto *D*^W^*−θ−ϕ* space. This newly defined reference frame reliably separates the open and closed states as was the case in the |*h*|*−θ*_cyl_ space. In the specific case of Wzc, this separation occurs along *θ* in addition to *D*^W^ (*SI Appendix* Fig. S2). Indeed, as will be shown in greater detail below, the mode of achieving the open state through a change along one, or both, of the two angular directions is a property that is dependent on the specific family of P-loop enzymes. Further, given the high correlation (*SI Appendix* Fig. S2) of *D*^W^ with both |*h*| (0.71) and *θ*_cyl_ (0.82), it serves as a direct proxy for the open (high values of *D*^W^) and closed (low values of *D*^W^) states.

### Structures of the KG branch of P-loop enzymes populate open and closed states

The KG dataset (Dataset S1) used in our analysis comprised 236 unique PDB entries for a total of 467 structures (see Materials and Methods for details) of the SIMIBI (named for <u>si</u>gnal recognition particle, <u>Mi</u>nD and <u>Bi</u>oD) and TRAFAC (named for <u>tra</u>nslation <u>fac</u>tors) classes (3). Individual chains from a single PDB entry were treated as separate entities to account for possible conformational variability at the active site especially with respect to key residues of the Walker-A and Walker-B motifs. Structures of the BY-kinase CDs, that belong to former class, and those of the kinase complement of the KG branch were excluded from the analyses since we have comprehensively analyzed their conformational dynamics previously and established the presence of global open and closed states (14–16).

The structures comprising the KG dataset were projected onto *D*^W^*−θ−ϕ* space and clustered using the Mean Shift (MS) algorithm (20). This protocol yielded 28 distinct clusters, of which only 7 contained more than 9 structures each accounting for 92% of the total number of structures in the dataset (430/467, *SI Appendix* Fig. S3). A decomposition of the three-dimensional MS-clustered distribution into 3 two-dimensional distributions is shown Fig. 3A. *D*^W^ values close to ∼4.3 Å indicate the presence of a hydrogen-bond between Ser_A_/Thr_A_ and Asp_B_. Therefore, all clusters with *D*^W^ centers (representing the most probable value along a given coordinate) below 4.8 Å e.g., green and dark-blue clusters (see *SI Appendix* Table S1 for cluster statistics), may be taken to represent closed states. Structures comprising the green cluster have higher values of *ϕ*, on average (97° vs 84°, see *SI Appendix* Table S1), compared to those of the dark-blue cluster, suggesting that Asp_B_ is positioned differently relative to *α*^P^ within these two clusters. The red, purple, pink, gray and black clusters are all characterized by high *D*^W^ values (center values more than 6 Å) and are indicative of open states. The differences in geometry for the open states is determined by variations along *ϕ* or *θ*, the implications of which will be discussed at length below.

The closed state structures comprising the green and dark-blue clusters are largely NTP•Mg^2+^ and NDP•Mg^2+^ bound, with Mg^2+^-bound species comprising 75% and 50% of the structures, respectively (Figs. 3B, C). In contrast, structures of the purple, black, pink, grey and red clusters, representing open states, are almost exclusively Mg^2+^-free (Figs. 3B, C). Indeed, only ∼5% of open state structures are Mg^2+^-bound, and these lie along the borders with the closed states. This suggests that this divalent cation cannot be optimally accommodated in the open state.

**Fig. 3.**
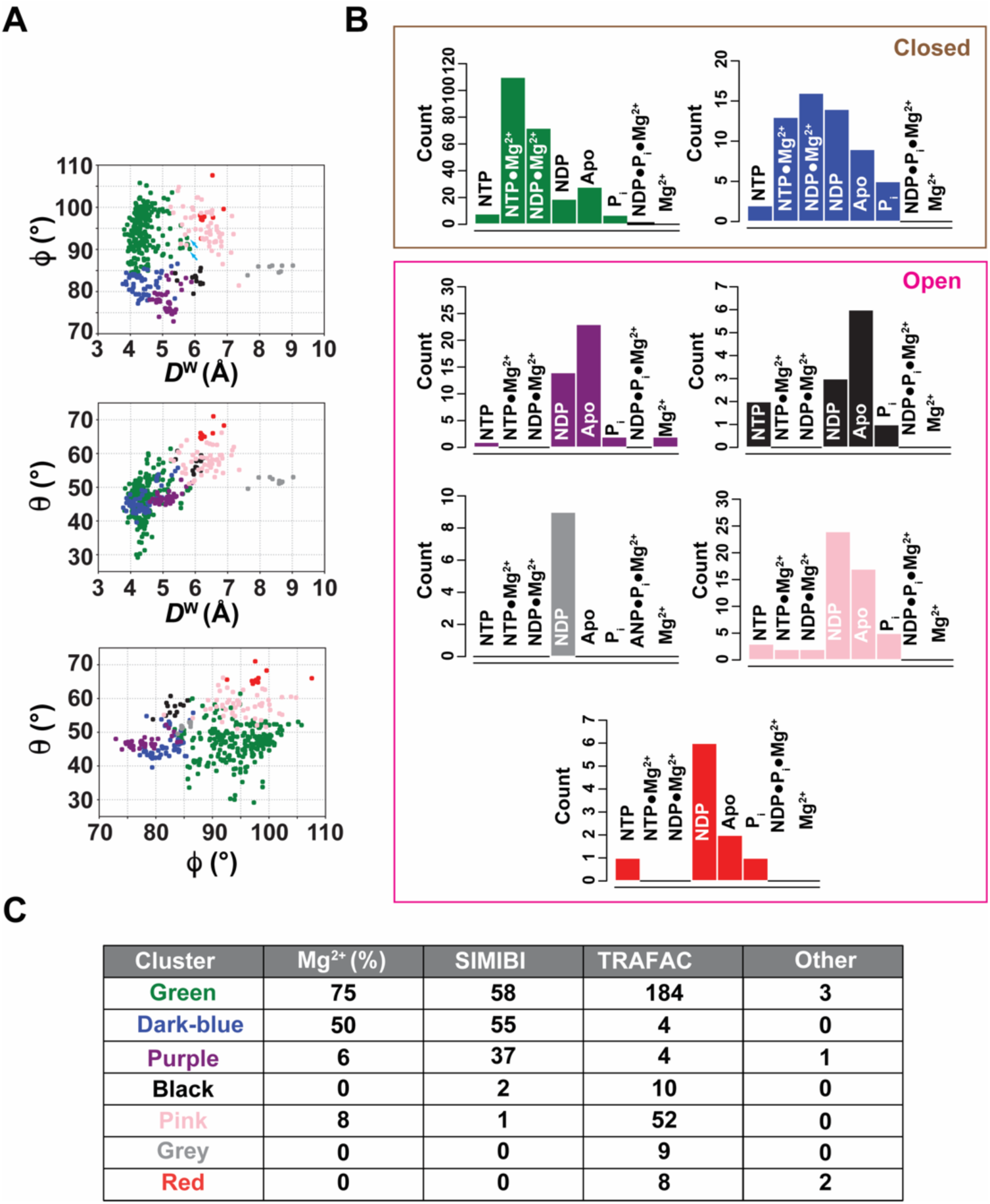
Analysis of KG structures in the spherical coordinate frame. **(A)** The 9 major KG clusters represented in *D*^W^*−ϕ−θ* space shown as three two-dimensional projections. Individual clusters are labeled using an arbitrary color scheme. Overall, the dark-blue and green clusters are characterized by low values of *D*^W^ and indicate closed states. The remaining clusters show large values for *D*^W^ indicative of open states. **(B)** Distribution of bound ligands depicted as histograms colored in accordance with the corresponding cluster in **(A)**. **(C)** Distribution of the SIMIBI or TRAFAC group members within the major clusters. Also shown are the percentages Mg^2+^-bound structures in each cluster. 6 structures that could not be definitively identified as being members of either the TRAFAC or SIMIBI groups from their PDB or UniProt annotations have been designated as “Other”.

MS clustering also appears to parse members of the TRAFAC and SIMIBI classes into different clusters, a pattern that is especially true for the open states. As shown in Fig. 3C, for the more significantly populated open clusters, TRAFAC structures are more likely to be in the pink cluster while SIMIBI structures are more likely to be in the purple cluster. For the clusters representing the closed states, the smaller dark-blue cluster is dominated by SIMIBI structures, while the significantly larger green cluster contains significant numbers of both SIMIBI and TRAFAC, with a somewhat larger number of the latter, in a proportion that is not widely different (SIMIBI: 24%, TRAFAC: 75%) from the relative populations of the two species in the KG dataset as a whole (SIMIBI: 36%, TRAFAC: 63%, Fig. 3C). These observations indicate that the open states more reliably cluster based on class i.e., SIMIBI or TRAFAC, suggesting that the geometry of the open states depend more strongly on the details of the global fold while the closed states are more similar across fold-families.

### Differences in interactions of the Walker-A Lys distinguishes open and closed states within the KG branch

As noted earlier for Wzc, the open state is characterized by an interaction of the sidechains of Lys_A_ and Asp_B_ (designated as the *α−*position) resulting in a configuration that is unsuitable to engage Mg^2+^. An additional interaction involving the Lys_A_ sidechain with the backbone of a residue immediately following Asp_B_ (Thr in Wzc; the *β−*position) further stabilizes the open state (Fig. 4A) (16). To test the universality of these interactions in the KG structures, we measured the distances of the N*ζ* atom of Lys_A_ with the C*γ* atom of Asp_B_ (*D^α^*) and the carbonyl O atom of the residue that follows (*D^β^*) (Fig. 4B). In the green cluster, which consists of closed state structures, both *D^α^* and *D^β^* tend towards larger values indicating that Lys_A_ has disengaged from the *α−*, *β−*positions on the Walker-B motif. A large majority of the structures of the purple, pink, black and gray clusters, i.e., those in the open states, show reduced *D^α^* and *D^β^* values (Fig. 4B) placing Lys_A_ within interacting distance of Asp_B_ (*α*-position, ∼3.5Å) and the residue that follows it (*β*-position, ∼2.7Å). The behavior seen for the dark-blue cluster, that is also representative of the closed state, is somewhat more complex. The *D^α^* and *D^β^* distributions for this cluster are bimodal, with one of the maxima (∼7.5 Å and 4.5 Å for *D^α^* and *D^β^* respectively) resembling that of the green cluster i.e., that expected for the closed states (Fig. 4B), while the other is characteristic (∼3.5 Å and ∼2.8 Å for *D^α^* and *D^β^* respectively) of the open states. It is notable that the “anomalous” structures that comprise the latter group are Mg^2+^-free (indeed the structures of the dark-blue cluster are only ∼50% Mg^2+^-bound, see Fig. 3C), however Lys_A_ is not sufficiently displaced relative to the Walker-B (see *SI Appendix* Fig. S4 for representative examples) suggesting that these structures are in intermediate conformations, that while globally closed (as reflected by the shorter *D*^W^ values), retain local features characteristic of open states.

**Fig. 4.**
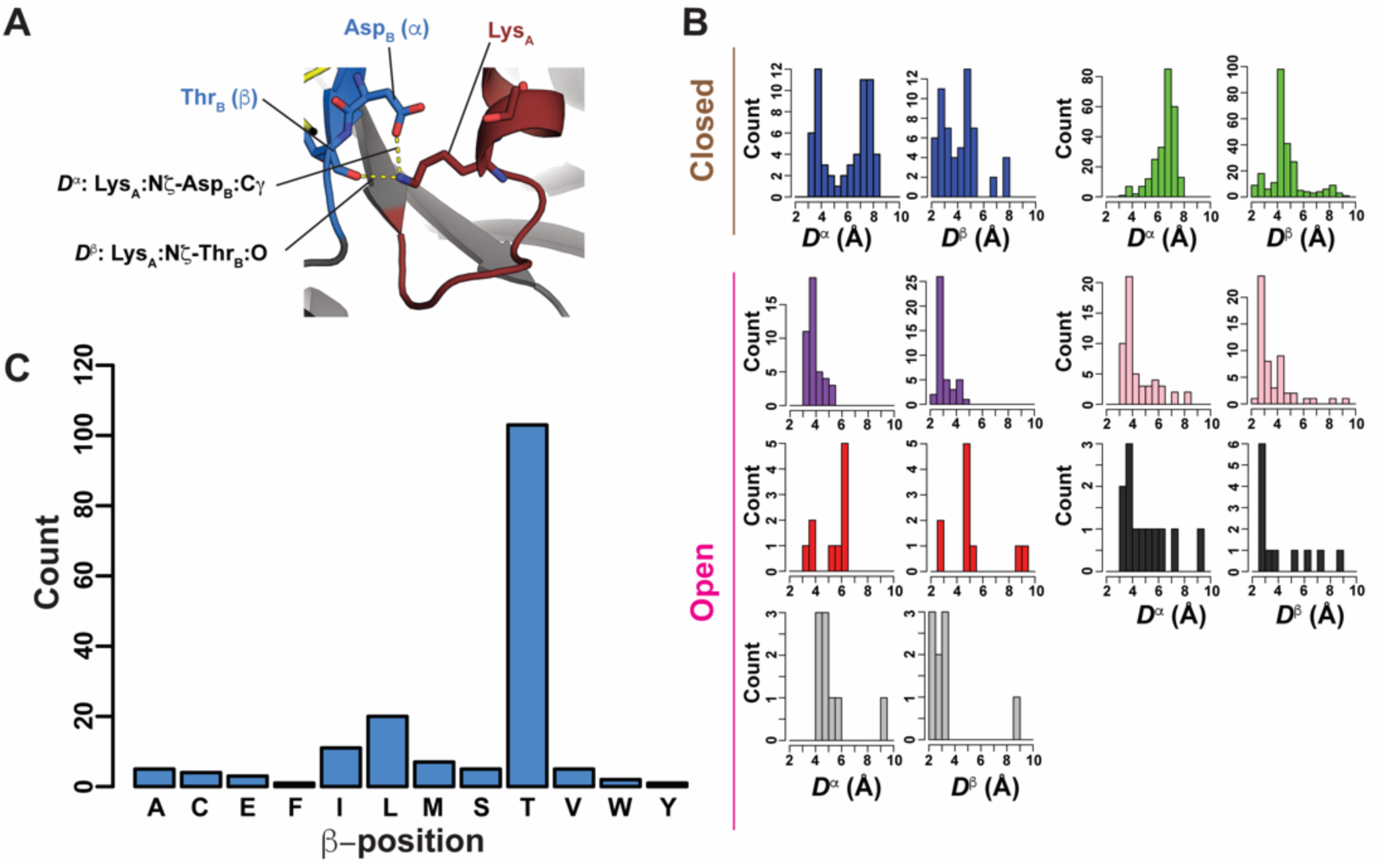
Local interactions between the Walker-A and Walker-B motifs. **(A)** In the open state of the catalytic core of Wzc, the interaction between of the sidechains of Asp_B_ and Lys_A_ are supplemented by an interaction of the latter with the backbone of a Thr (Thr_B_) that immediately follows Asp_B_ (yellow dashes). For analyses of the KG structures, the residues corresponding to Asp_B_ and Thr_B_ in Wzc are designated as the *α−* and *β−*positions, respectively. *D^α^* and *D^β^* measure the distances between the Lys_A_ N*ζ* atom and the *α−* C*γ* (or C*δ* when this position is a Glu) and *β−* carbonyl oxygen, respectively. **(B)** Distribution of *D^α^* and *D^β^* distances in the various clusters (colored using the same scheme as in Fig. 3) represented using 0.5 Å bins. **(C)** Distribution of amino-acid type at the *β−*position over the KG dataset.

As in the case of Wzc, the residues at the *α−*position for members of the SIMIBI and TRAFAC classes is either an Asp or a Glu. This is not true for several families of the kinase complement of the KG branch, as discussed below. An analysis of residue-types at the *β−*positions within the various clusters provides some further interesting insights. The most commonly occurring residue at this position is a Thr, as is the case in *E. coli* Wzc where we first identified its importance in stabilizing the open state (16), with Leu being a distant second (Fig. 4C). In addition to an interaction of its backbone with the Lys_A_ sidechain, the residue at the *β*-position utilizes its sidechain in a variety of ways to stabilize the open state. The precise mode of interaction depends on the tertiary structure of the protein in question (see *SI Appendix* Fig. S5 for some representative examples).

### A residue on *α*^P^ places geometric constraints on the open state in KG structures

As noted earlier, there is considerably more structural diversity in the open states relative to the closed states. Comparing, for example, the dark-blue and purple clusters that comprise mostly of closed and open SIMIBI structures, respectively, specific trends can be seen. With an increase in *D*^W^ there is a concurrent decrease in *ϕ* (see the top panel in Fig. 3A) implying an upward motion of the Walker-B containing *β*3 relative to *α*^P^ (Fig. 5A). In contrast, for the corresponding TRAFAC structures (compare the green and pink clusters where these structures dominate in Fig. 3C), opening occurs with almost no change in *ϕ* but a significant increase in *θ* (compare the top and middle panels of Fig. 3A) due to a clockwise rotation of *β*3 about *α*^P^ (Fig. 5A). The black and gray open clusters (Fig. 3A), also largely consisting of TRAFAC structures, also show reduction in the *ϕ* angle compared to the green cluster (see the top panel of Fig. 3A) in similar fashion as the structures of the SIMIBI family. We wondered whether any specific patterns that predict the overall mode of opening could be ascertained. We found that in cases where opening occurred largely through a change in *ϕ* (purple, black and grey clusters), the 4^th^ position (*K*_i+4_) relative to Lys_A_ on *α*^P^ is generally a polar residue (Ser or Thr) or a Phe (right panel of Fig. 5B). In contrast, for the cases where opening involves substantial changes in *θ* (pink and red clusters) this position almost always contains a bulky hydrophobic residue (mostly Leu or Ile; left panel of Fig. 5B). Thus, based on this analysis it appears that the *K*_i+4_ position correlates with the mode of opening. When opening accompanies changes in *ϕ*, the corresponding *K*_i+4_ residue, can take part in specific interactions with Asp_B_ either through the formation of hydrogen-bonds when a Ser/Thr is present, or through anion-*π* contacts (21) (22) in the presence of the aromatic ring of Phe. In cases, where such an interaction is not possible e.g., in the presence of an aliphatic sidechain, opening may involve changes in *θ*. Representative examples are shown in *SI Appendix* Fig. S6.

**Fig. 5.**
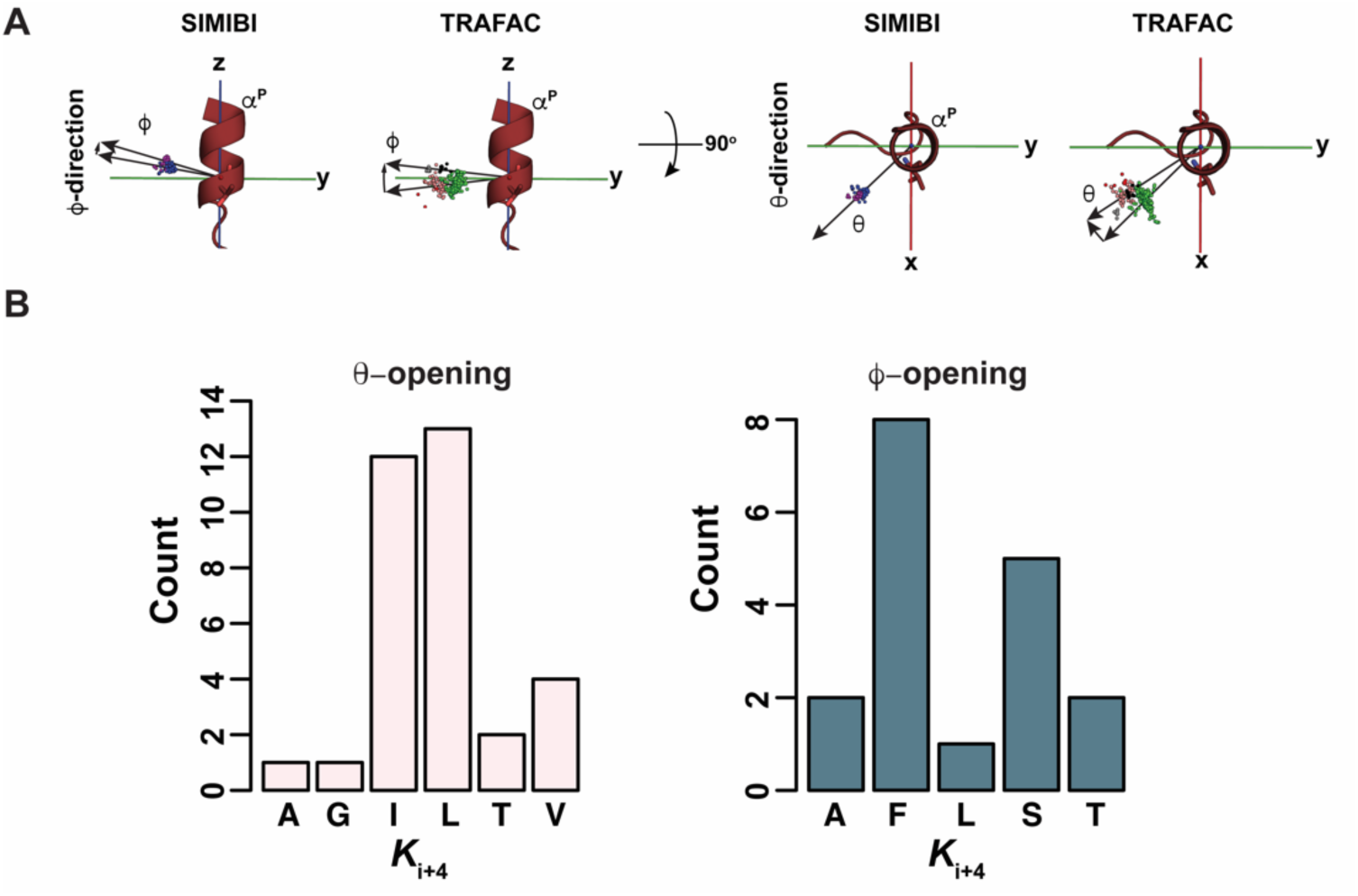
Importance of the *K*_i+4_ residue in stabilizing the open state. **(A)** Positions of the Asp_B_ C*α* atoms for the structures of the KG dataset are represented as spheres colored in accordance with the cluster they belong to. The *α*^P^ helix is shown in brown for reference. The left panels show the yz-projections illustrating changes in *ϕ* upon undergoing global opening transitions for the structures belonging to the SIMIBI (left) and TRAFAC (right) classes. The right panels depict the xy-projections and the corresponding changes in *θ*. **(B)** Distribution of residue types at the *K*_i+4_ is shown for the clusters that open largely along *θ* (left) and those that open along *ϕ* (right).

### The presence of open and closed states is also a feature of ASCE structures

In order ascertain the generality of the features discovered in the KG structures to a broader class of P-loop enzymes, we extended our analyses to the members of the ASCE group. The ASCE dataset used in the analysis comprised of 296 unique PDB entries corresponding to 734 total structures (Dataset S2). These structures were transformed into the spherical coordinate system (Fig. 2) and clustered using the MS algorithm as before. The procedure yielded 28 clusters of which only 11 contained more than 10 structures each accounting for 94% (692/734) of the total structures in the dataset (*SI Appendix* Fig. S7). The dark-blue and green clusters are characterized by low *D*^W^ values (< 4.8 Å, see *SI Appendix* Table S2) suggesting closed states (Fig. 6A). The other 9 clusters (red, magenta, grey, yellow, pink, black, brown, lime and purple) with larger *D*^W^ values (see *SI Appendix* Table S2) are indicative of open states. As in the case of the KG structures, the clustering procedure also parses Mg^2+^-bound and Mg^2+^-free states (Fig. 6B) together with the constituent fold families within the ASCE branch (Fig. 6C).

**Fig. 6.**
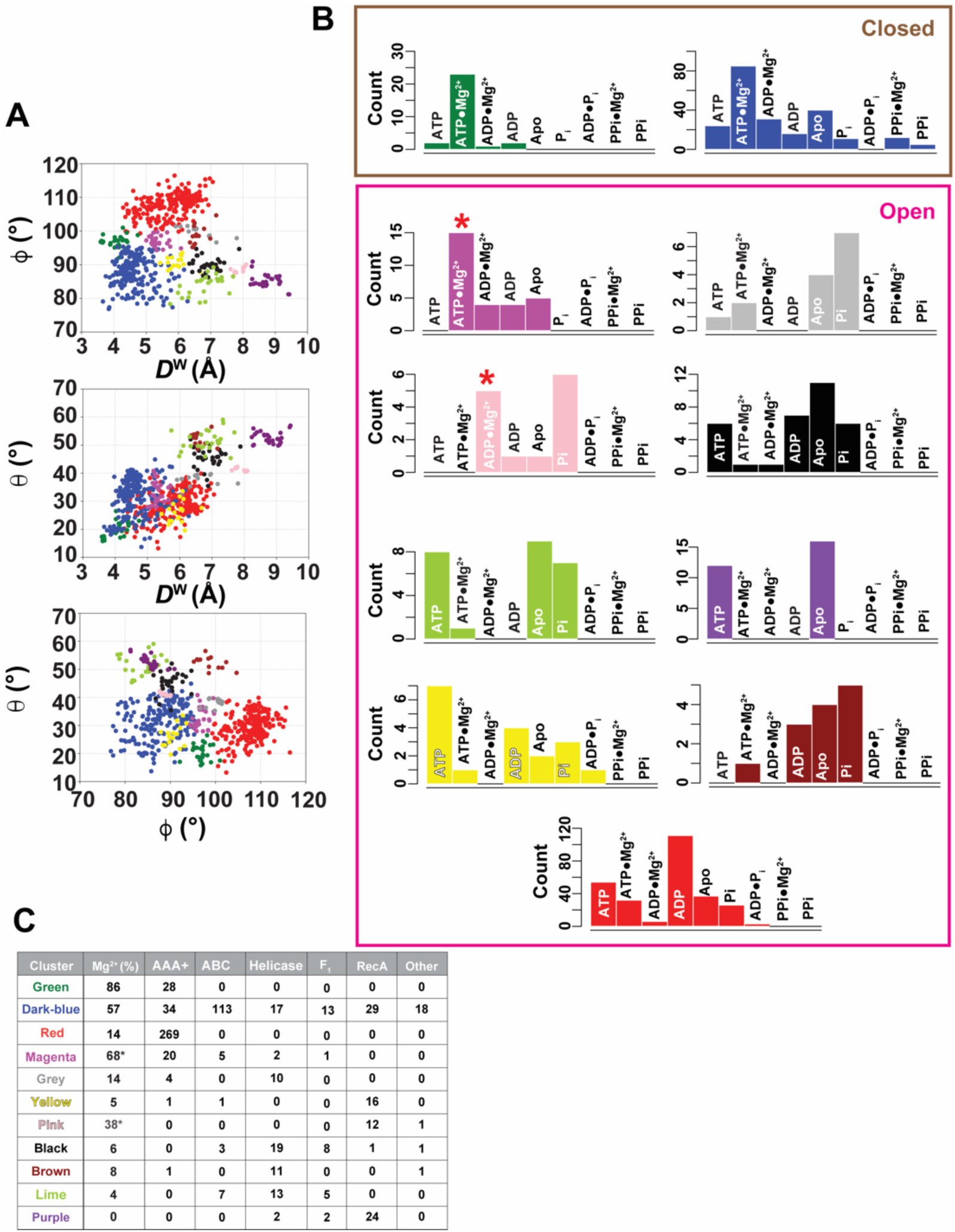
Analysis of ASCE structures in the spherical coordinate frame. (**A**) Projection of the major clusters onto the spherical coordinate frame as in Fig. 3. The dark-blue and green clusters have low *D*^W^ values and represent closed state structures; the reminder of the clusters consist of open state structures with high values of *D*^W^. **(B)** Distribution of liganded states of the different clusters colored as in (A). **(C)** Distribution of specific families from the ASCE branch between the various clusters. Also shown are the percentages Mg^2+^-bound structures in each cluster. 21 structures that could not be definitively identified as being members of any of the represented families from their PDB or UniProt annotations have been designated as Other. The “anomalous” magenta and pink clusters designated as open but have substantial proportions Mg^2+^-bound structures (a plurality for the magenta cluster) are indicated by ‘*’.

The dark-blue cluster, the largest cluster corresponding to closed states, contains members from all the ASCE sub-classes, namely the AAA+, ABC transporter, helicase, F_1_-ATPase and RecA families, suggesting the similarities in the global properties of the closed conformation as noted for the KG structures above. The structures comprising this cluster are predominantly Mg^2+^-bound (57%). The green cluster, which also contains closed state structures, is comprised exclusively of AAA+ members that are also predominantly Mg^2+^-bound (87%, Fig. 6C). In contrast, the largest grouping of open states, the red cluster that consists exclusively of AAA+ structures that are predominantly Mg^2+^-free (14% bound). Indeed, the clusters corresponding to the open states are largely dominated by members of a single family, e.g., the yellow, pink, and purple clusters are predominantly RecA family members, while the black, brown, and grey clusters are dominated by helicases. As in the case of the KG structures, the open states are predominantly Mg^2+^-free except the magenta cluster (mainly AAA+), and to some extent the pink cluster (RecA), that contain a substantial proportion of Mg^2+^-bound structures. The reasons for these apparent exceptions are discussed in greater detail below. Nevertheless, consistent with the trends observed for the KG structures, members of the ASCE branch also populate globally open and closed states, where the Asp_B_ and Ser_A_/Thr_A_ interaction is established in the latter and broken in the former that are less likely to be engaged to Mg^2+^.

As mentioned in the previous paragraph, the structures of the magenta cluster that represent open states, are frequently Mg^2+^ -bound (68%; Fig. 6C). Inspection of these “anomalous” structures confirms the presence of the divalent cation through the presence of representative electron density (*SI Appendix* Fig. S8). However, in these structures, Asp_B_ is clearly displaced from Ser_A_/Thr_A_, and Lys_A_ has moved away from the nucleotide, all features characteristic of open states. It therefore appears that the Mg^2+^-bound structures of the magenta cluster represent conformations that are primed to eject the Mg^2+^ ion and likely represent intermediates on the pathway to full opening. Indeed, the *D*^W^ center for the magenta cluster (5.2 Å; see *SI Appendix* Table S2) is only marginally higher than our cutoff (4.8 Å) for a closed state.

A significant fraction of the structures of the pink cluster, that largely comprises of RecA family members, is also Mg^2+^-bound despite being classified as open (Fig. 6C). Comparison of two sets of structures of the RecA family member TrwB (23), one from the pink cluster bound to ADP•Mg^2+^ and another from the purple cluster bound to the GTP analog, GMP-PNP, reveals a shift of Asp_B_-bearing *β*3 strand towards *α*^P^ in the latter. However, the Asp_B_ sidechain is modeled such that it faces away from Ser_A_ resulting in a large *D*^W^ (*SI Appendix* Fig. S9). Correspondingly, the structure (lower panel in *SI Appendix* Fig. S9) is also globally shifted in the same direction as *β*3. These differences are representative of those between open and closed states suggesting that the Mg^2+^-bound members of the pink cluster contain features characteristic of the closed states.

### The additional *β*-strand in ASCE members is structurally equivalent to the degraded Walker-B in SKs

As discussed earlier for the KG branch, the *α−* and *β−*positions on the Walker-B motif play critical roles in stabilizing the open state (see Fig. 4). The SKs that are part of the KG branch, as mentioned above, contain degraded Walker-B motifs that lack the characteristic Asp_B_ (or Glu_B_) at the *α*-position. The corresponding role in Mg^2+^-coordination is replace by the first Asp of a so-called DXD (where X is any residue) motif housed on a strand (*β*2) immediately following *α*^P^ (Fig. 7A, left panel). We have previously shown that despite the absence of a characteristic acidic residue at the *α−*position, it and the corresponding *β−*position, also stabilize the open state of SKs (16). A DXD motif (referred to as the Walker-A’ motif in the BY-kinase literature) (24) is found at a similar position in Wzc, that also contains an intact Walker-B with the constituent Asp_B_ (Fig. 7A, middle panel). In ASCE ATPases, the additional *β*-strand (*β*4) is sandwiched between *β*1 and *β*3 at an equivalent spatial position as the degraded Walker-B in SKs. The corresponding *β*3 strand carrying the Walker-B motif is spatially equivalent to the *β*2 strand that houses the DXD motif in the SKs (Fig. 7A, right panel) suggesting a degree of structural equivalence between Walker-B motif in ASCE ATPases and the DXD motif in the SKs, and between *β*4 in ASCE ATPases and *β*3 that houses the degraded Walker-B motif SKs. Thus, it is tempting to speculate the *α−* and *β−*positions of the degraded Walker-B motif in SKs are encoded within *β*4 in the ASCE group members.

**Fig. 7.**
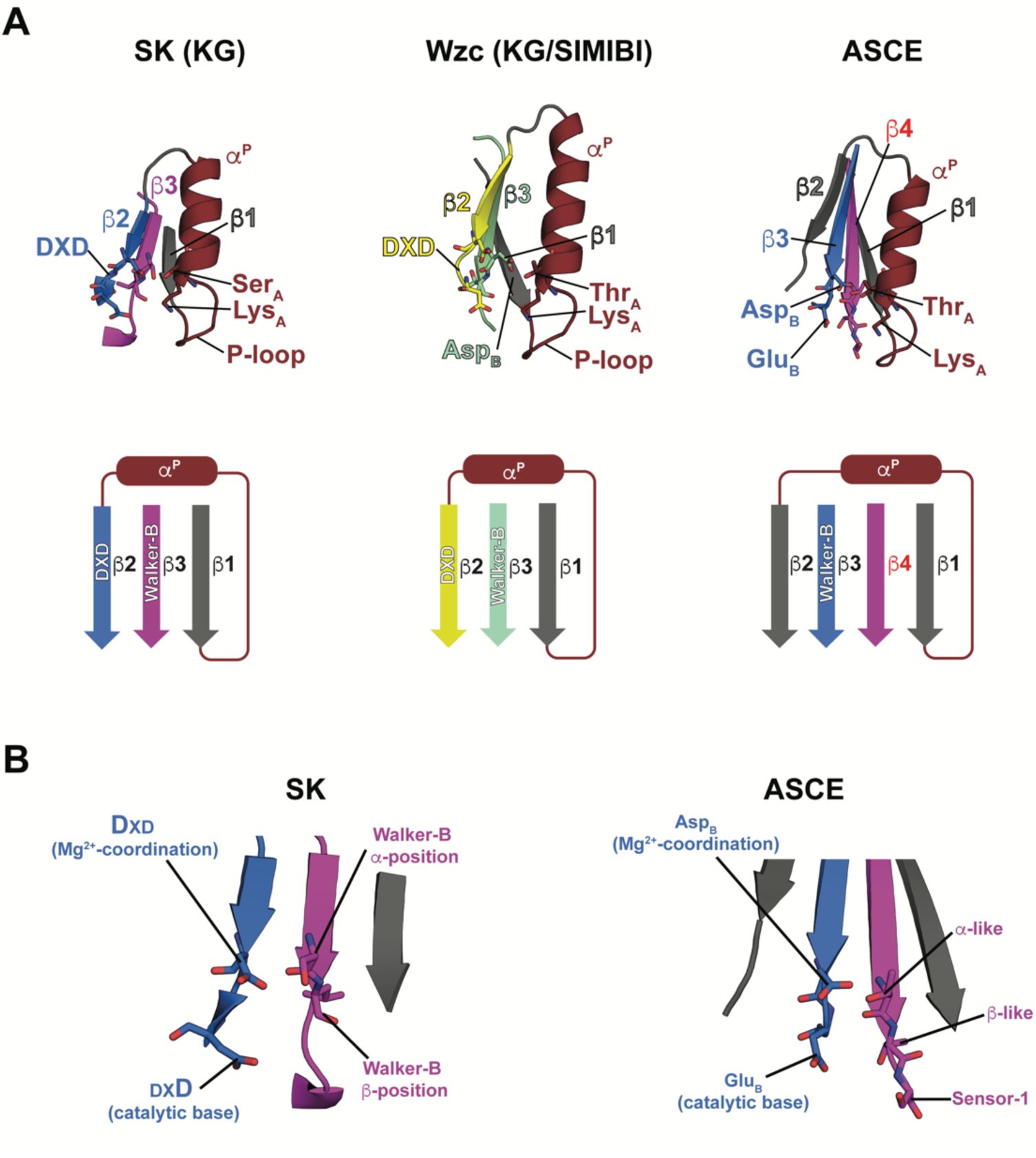
Comparison of spatial arrangement of key secondary structure elements in P-loop enzymes. **(A)** Arrangement of key structural elements are shown for SK (left, KG branch kinase), Wzc (middle, tyrosine kinase classified as a SIMIBI GTPase) and a representative member of the ASCE ATPase group. The corresponding topology is shown schematically in the bottom panels. The first *β*-strand (*β*1, grey) continues into the P-loop and the *α*^P^ helix (brown), and the second strand (*β*2) that houses the DXD motif in Wzc (yellow) and SK (blue) lie in a continuous sequence. The proximal *β*-strand (*β*3) contains the degraded Walker-B motif in SK (magenta), the intact Walker-B motifs for Wzc (mint-green) and ASCE (blue; the catalytic Glu_B_ immediately follows Asp_B_). The extra strand (*β*4, magenta) inserts between *β*1 and *β*3 in ASCE ATPases. **(B)** Comparison of the arrangement of key “catalytic” residues in SKs (left) and ASCE ATPases (right). Functionally equivalent residues are indicated. The putative *α−* and *β−* equivalents of the degraded Walker-B in SKs (on *β*3, magenta) lie on the additional strand (*β*4, magenta) in ASCE ATPases. Also indicated is the polar Sensor-1 motif that plays a key role in hydrolysis in ASCE ATPases (39).

To confirm the equivalence between structural elements on the SKs and ASCE members, we constructed a reference set of 100 structures drawn randomly from the closed states observed in our previously described enhanced sampling simulations of the ATP•Mg^2+^ complexes of SK (16) and Wzc (14). These structures were transformed into spherical coordinates and the C*α* atoms of the first Asp of the corresponding DXD motifs and the Walker-B *α*-positions (Asp in Wzc and Ser in *M. tuberculosis* SK) were projected onto the xy-plane (Fig. 8A, left panel). Relative to its counterpart in Wzc (yellow), the DXD motif of SK (blue) is displaced in a clockwise fashion by approximately 10° (in *θ*) with respect to the helical axis of *α*^P^. This has the additional effect of positioning the first Asp of the DXD motif of SK closer to Thr_A_ compared the corresponding element in Wzc. The *α−*position of the degraded Walker-B in SK (magenta spheres) is also shifted in the same direction relative to its counterpart in Wzc (Asp_B_, mint-green spheres) by about 13°. The distribution of the corresponding *θ* angles illustrate the overall shifts (Fig. 8B, top panels). To test the proposed equivalence with structural elements in the ASCE ATPases, we constructed an equivalent set of projections using the 357 structures of AAA+ ATPases that comprise a significant proportion (∼49%) of the ASCE structures used in the present study. This was done to obtain a homogenous dataset unobscured by structural differences between different ASCE families. The maxima of the distributions corresponding to Asp_B_ and the putative *α*-position on *β*4 in the AAA+ structures coincide with those of the first Asp of the DXD motif and the *α*-position on the degraded Walker-B motif in SKs, respectively, suggesting their overall equivalence (Fig. 8B, bottom panels). Visual inspection of other ASCE families confirms that this structural equivalence holds for all ASCE ATPases. Note that the distribution of Asp_B_ positions in the AAA+ family shows a tail that also overlaps with the maximum of the first DXD Asp of Wzc (indicated by the arrow in the bottom panel of Fig. 8B) further highlighting the equivalence.

**Fig. 8.**
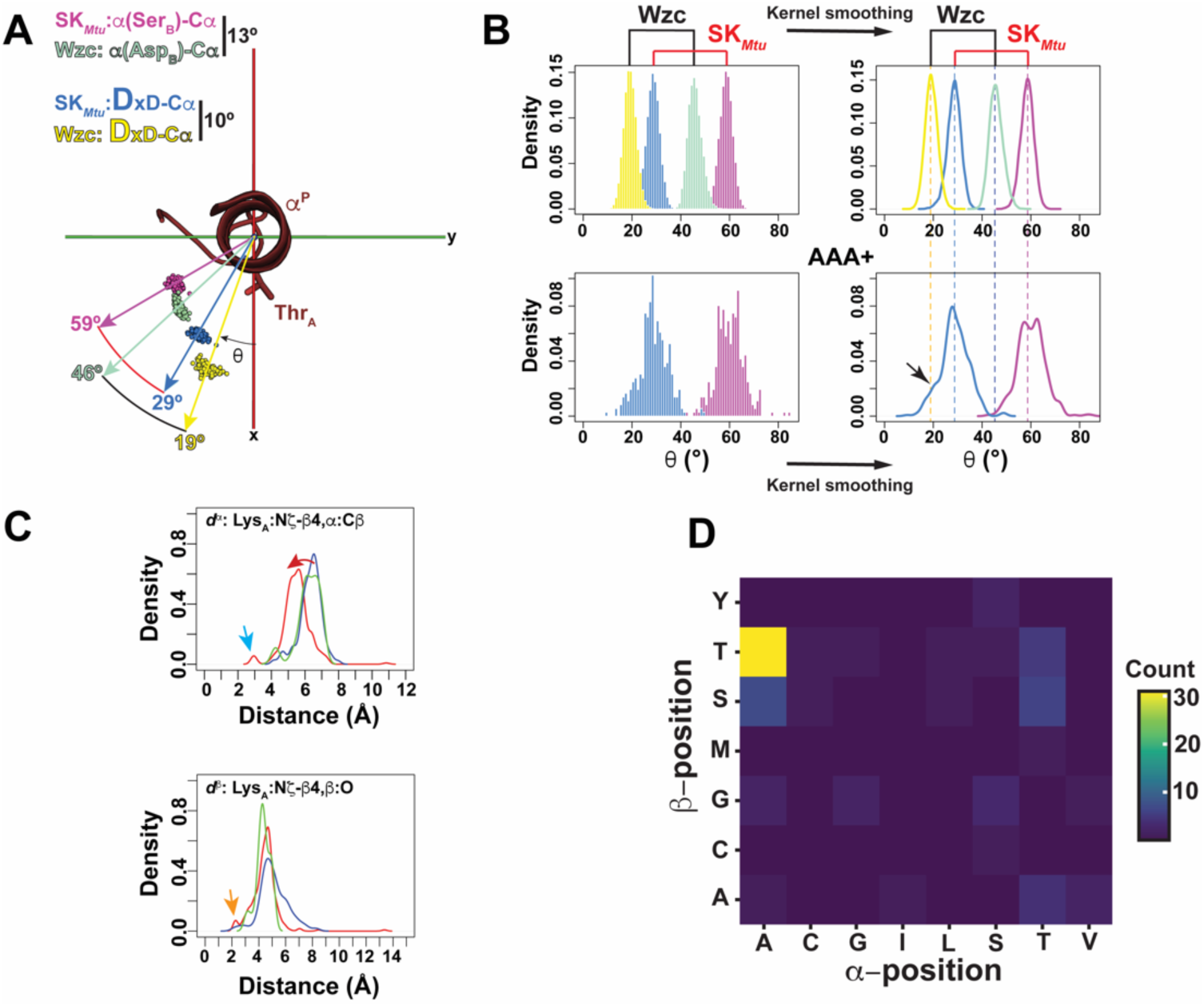
Comparison of key structural motifs of ASCE ATPases and SKs. **(A)** 100 structures of *M. tuberculosis* SK (SK*_Mtu_*) (16) and *E. coli* Wzc (14) drawn from the closed states sampled in the enhanced sampling simulations of their corresponding ATP•Mg^2+^ complexes transformed into the spherical coordinate system and projected onto the xy-plane. The C*α* atoms of the first Asp of the DXD motif (shown in larger font; SK*_Mtu_*: blue; Wzc: yellow) and the *α−*positions (SK*_Mtu_*: Ser_B_, magenta; Wzc: Asp_B_, mint-green) are shown as spheres indicating the subtended *θ* angles. The Wzc the structures were extracted from simulations on a specific 4-glycine mutant that only samples closed states when complexed with ATP•Mg^2+^ (14). **(B)** Distribution of C*α* positions from (A) over the entire trajectory shown as 1° binned normalized histograms on the top left panels. The top right panels show kernel density estimates of the corresponding histograms with the dashed vertical lines marking the most probable values of the Wzc and SK*_Mtu_* distributions. The corresponding distributions for the AAA+ ATPases for all crystal structures used in the analysis are shown in the bottom panels. To highlight their equivalence, the Asp_B_ distribution for the AAA+ structures is colored blue (as for the DXD in SK*_Mtu_*) and the additional *α−*position equivalent (on the addition strand, *β*4) is colored magenta (as for the degraded Walker-B motif in SK*_Mtu_*). **(C)** Normalized estimated densities after kernel smoothing of the distances between Lys_A_ and the *α−* and *β−*positions on *β*4 in AAA+ ATPases are shown. The green and red clusters (see Fig. 6) correspond to closed and open states, respectively, of the AAA+ ATPases. Also shown for comparison are the corresponding distributions for the dark-blue cluster that includes closed state structures from all the ASCE ATPase families. The red arrow indicates the shift towards shorter values of *d^α^* upon opening. The orange and cyan arrows indicate specific contacts highlighted in *SI Appendix* Fig. S9. **(D)** Pairwise frequency of occurrences of amino acid types at the *α−* and *β−* positions in AAA+ ATPases.

As noted earlier, in the KG group members, the *α−* and *β−* positions on the Walker-B motif play key roles in stabilizing the open states (Fig. 4). We examined whether the *α−* and *β−*equivalents on *β*4 play a similar role in the ASCE ATPases. We defined a set of distances between Lys_A_ N*ζ* and the sidechain C*β* of the *α*-position (*d^α^*, corresponds to with *D^α^* in Fig. 4; however, note that these are not identical measures) and the carbonyl oxygen of the *β*-position (*d^β^*; to imply correspondence with *D^β^* in Fig. 4) for AAA+ ATPases in the open (the red cluster in Fig. 6) and closed states (green cluster in Fig. 6; the dark-blue cluster that also contains AAA+ structures is also shown for reference). The maximum of the *d^α^* distribution suggests shorter values in the open (red) compared to the closed (dark-blue, green) states, as expected. However, the corresponding distances are larger than that expected for a direct interaction, that is only occasionally seen (top panel of Fig. 8C, cyan arrow). Shifts towards shorter distances for the *d^β^* distributions upon comparing open and closed states are less evident though direct contact is also occasionally seen (bottom panel of Fig. 8C, orange arrow) in the open states. These results suggest, though less clearly so, a similar role for *α−*equivalent on the ASCE *β*4 in maintaining the open states like their Walker-B counterparts (intact or degraded as in SKs) within the KG group. The role of the *β−*equivalent is less apparent. Representative structures highlighting these features are shown in *SI Appendix* Fig. S10. Additionally, the distribution of residue-types at the putative *α−* and *β−*positions on *β*4 for the ASCE class, indicates that AT is the most common pair (Fig. 8D). Note that the consensus sequence at these positions for SKs is *ϕ*T where is *ϕ* is a hydrophobic residue, most commonly an Ala (a non-standard SL is seen for *M. tuberculosis* SK) (4).

### The *K*_i+4_ residue stabilizes the open state in non-AAA+ ASCE ATPases

We had previously described a role for the *K*_i+4_ residue on *α*^P^ in defining the mode of opening within the KG structures depending on the presence/absence of stabilizing interactions with Asp_B_ (see *SI Appendix* Fig. S6). We wondered whether the corresponding residue also plays similar role in the ASCE ATPases. An inspection of the composition of the open clusters indicates that the black, brown, purple, lime, pink, and grey clusters, that are sparse in their AAA+ content, have enhanced *θ* values in addition to the characteristic high *D*^W^ values relative to the closed states middle panel of Fig. 6A). Given the relatively small number of structures in these clusters, we visually inspected the orientations of Lys_A_ and the *α−* and *β−*positions on *β*4. The lime, black, brown, and purple clusters that, on average, have similar *θ* values (middle panel of Fig. 6A), display similar features in the open state (see *SI Appendix* Fig. S11 for representative structures). In the open state structures of members of the helicase, F_1_-ATPase, ABC transporter, and RecA families, the Lys_A_ sidechain lies proximal to the sidechains of Asp_B_ and Glu_B_, and in many cases forms salt-bridges with one or both (*SI Appendix* Fig. S11). The *K*_i+4_ residue, in conjunction with the residue at the *α−*position on *β*4, provide a platform for the aliphatic part of the Lys_A_ sidechain to maintain it in an extended conformation. A direct interaction between the *α−* and *K*_i+4_ sidechains, somewhat reminiscent of that in that seen for the KG branch (*SI Appendix* Fig. S6), is seen in virtually all open state structures except in RecA (purple and pink clusters) that contains Gly at the *α−*position. Representative examples are shown in *SI Appendix* Fig. S11. Indeed, the interdependence between these two positions is evident from the several cases where residue-types for the *K*_i+4_/*α* pairs are swapped (i.e., L/T to T/L or A/I to I/A indicated by a ‘*’ or ‘#’, respectively in the bottom panel of Fig. 9A) presumably to maintain a constant distance between the two sets of sidechains. A similar analysis using representative sequences also produces comparable results (see *SI Appendix* Fig. S12). In contrast, an analysis of the structures of the red cluster (Fig. 6) that comprises exclusively of members of the AAA+ family, A/A is the dominant *K*_i+4_/*α* pair (top panel in Fig. 9A). It is also significant that members of AAA+ family achieve their open states through changes in both *ϕ* and *θ* (compare the green and red clusters in Fig. 6A) in contrast to the other ASCE ATPases for which opening principally involves changes in *θ*. Indeed, the values of *D*^W^ show relatively strong correlation with both the *θ* (0.44) and *ϕ* (0.63) angles for the AAA+ ATPases in contrast to the ABC transporters, helicases, F_1_-ATPases and RecA family members, where there is substantial correlation only with *θ* (0.72; see Fig. 9B). In AAA+ ATPases since both the *α−* and *K*_i+4_ positions are occupied by small amino acid residues, reducing the possibility of a direct interaction, allows these proteins to open by varying both *ϕ* and *θ* i.e., through a forward translation and a clockwise rotation of Asp_B_ relative to *α*^P^ (shown schematically in the left panel in Fig. 9C). For the remaining ASCE members, the *α−* and *K*_i+4_ positions involve pairs of residues that in combination generate a more significant excluded volume resulting in a steric barrier that restricts the Walker-B motif to a fixed distance from *α*^P^. Opening can therefore only occur along the *θ* (right panel of Fig. 9C).

**Fig. 9.**
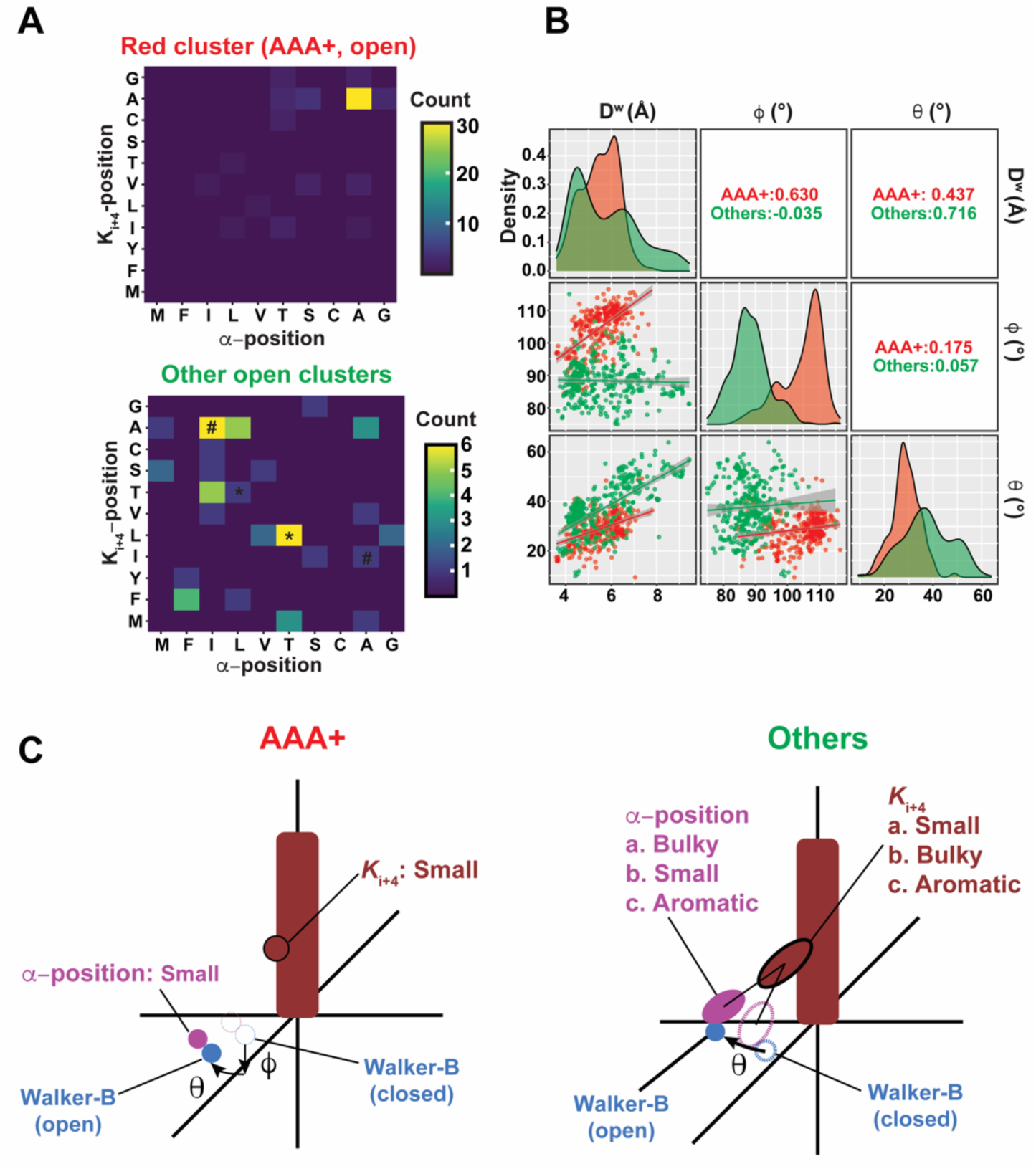
Dependence of the mode of opening on the *K*_i+4_ position in ASCE ATPases. **(A)** Pairwise occurrence of amino acid residues at the *α−* and *K*_i+4_ positions in the open state clusters: red cluster (exclusively AAA+ ATPases, top panel) and all other clusters (bottom panel). The ‘*’ and ‘#’ (bottom panel) indicate direct swaps of residue types between the *α−* and *K*_i+4_ positions. **(B)** Correlation between *D*^W^, *ϕ* and *θ* for AAA+ and non-AAA+ ASCE ATPases (others). The diagonal elements show *D*^W^, *ϕ*, and *θ* distributions for the AAA+ ATPases (red) and the others (green). The lower triangular matrix elements show the pairwise plots along with the corresponding linear regression lines (AAA+ in red, others in green) with shaded regions indicating the corresponding confidence bounds. The upper triangular matrix elements show the pairwise Pearson correlation coefficients for each case. **(C)** In AAA+ ATPases (left) both the *α−* and *K*_i+4_ positions contain small residues preventing their direct interaction. Opening can therefore occur through changes in both *ϕ* and *θ* i.e., through a downward shift and a clockwise rotation of the Walker-B motif with respect to *α*^P^. This is reflected in a positive correlation *D*^W^ with both *θ* (a somewhat larger correlation) and *ϕ* in (B). For the non-AAA+ ASCE ATPases, the nature of residues at the *α−* and *K*_i+4_ positions are such that they prevent downwards displacement of the Walker-B motif and opening can occur only through an increase in *θ* as reflected in the absence of correlation of *D*^W^ with *ϕ*, and its enhanced correlation with *θ* as shown in (B).

## Discussion

In this study we analyzed a large library (1201 structures from 532 distinct PDB entries) of P-loop enzyme structures representing both their constituent KG and ASCE branches (3, 4) as proxies for the sampled conformational landscape. To facilitate the analyses, we developed a generalized frame of reference to relate global conformational states to those of the conserved residues of the Walker-A and Walker-B motifs. Our analyses show that all P-loop enzymes populate global states, broadly classified as open or closed, in a manner that is strongly correlated with the local interactions at the catalytic site involving the Walker motifs. In the closed states, the conserved Asp_B_ is in contact with Ser_A_/Thr_A_, locked in a configuration necessary to coordinate the Mg^2+^ ion essential for catalysis. This key interaction is broken in the open states that are therefore incompatible with chemistry. The altered interactions at the catalytic site, a corresponding shift in the *β*3-strand that harbors the Walker-B motif relative to *α*^P^ that houses the Walker-A motif and the coupling of these local changes to the overall fold distinguishes the open and closed states. The closed states across multiple families are similar in overall geometry, contrasting the open states that are highly variable and dependent on the family or class of the P-loop enzyme. We suspect that this results from the fact that the geometric constraints on the closed state/s are imposed by chemistry, specifically by the ability to form an appropriate Michaelis complex to enable efficient transfer of the *γ*-phosphate of the NTP to a suitable target. The conformations of key elements that comprise this Michaelis complex are largely invariant among various P-loop enzyme families ensuring similar closed states. These arguments mirror those that have been invoked previously in describing the similarity of the active, and the “plasticity” of the inactive states of eukaryotic protein kinases (25).

Despite their diversity, some broad commonalities are also identified within the open states. For members of the KG branch, Lys_A_ establishes a direct interaction with sidechain of Asp_B_ (a position designated as *α*) and the backbone of the Walker-B residue that immediately follows it (*β*). However, though the *β*-sidechain stabilizes the open state, the mode by which this is achieved varies among the various constituent families. In the ASCE ATPases, Lys_A_ is also found to be proximal to the spatial equivalents of the *α−* and *β−*positions that are now hosted on the additional strand (*β*4), rather than *β*3 that hosts the Walker-B motif, also establishing direct interactions with Asp_B_ and catalytic Glu_B_ of the latter motif in some cases. Indeed, as for the KG group, while the *α−*, and to some extent the, *β−*sidechains for the ASCE ATPases also appear important within the open state, the mode that these are deployed is divergent among the various families. Similar variability in the mode of interaction is seen for the newly identified *K*_i+4_ residue on *α*^P^, that also stabilizes the open state, but does so in a variety of ways of depending on the type of P-loop enzyme.

Thus, there are primary features that are essentially identical for all P-loop enzymes. These include the alignment of the Asp_B_ and Ser_A_/Thr_A_ sidechains in a configuration suitable to coordinate Mg^2+^ in the closed state, and an inactive conformation of Lys_A_ that is diagnostic of the open state. In addition, family-specific diversity in the open state is obtained through secondary features involving differential use of the sidechains of the *α−*, *β−* and *K*_i+4_ residues that couple to the global fold of the specific P-loop family. Nevertheless, the defining feature of the closed state appears to lie in its ability to optimally coordinate Mg^2+^ and therefore, its associated NTP. It is notable that a given NTP, a substrate for a P-loop enzyme, has a significantly higher affinity for Mg^2+^ than the corresponding NDP (∼20-fold higher for ATP compared to ADP) (26), a reaction product. This difference, together with the NTP/NDP ratio, likely plays a critical role in facilitating nucleotide exchange and function in all P-loop enzymes. We argue that a primary driver in the activation of a P-loop enzyme lies in its ability to efficiently engage the Mg^2+^ ion, a feature that is only possible in the closed state, and not in the open state. However, the precise sequence of events, whether the open to closed transitions are induced by Mg^2+^, whether the local changes drive the global changes or vice versa, cannot be determined by the present analysis, and these may indeed be system dependent. For example, the F_1_-ATPase undergoes nucleotide cycling in its *β*-subunit subject to the *γ*-stalk rotation which in turn is driven by the proton motive force. The *β*-subunit transits between open and closed states driven by the rotation of the stalk domain (27) accompanied by catalytic site rearrangements like those discussed here (28). In Wzc, the closed state is stabilized by oligomer formation, aligning the catalytic site elements for the optimal coordination of Mg^2+^•ATP (14). In both these cases, the active closed state is formed but through somewhat through disparate means. Other studies have highlighted the role of Mg^2+^ in facilitating nucleotide exchange e.g., the engagement of a guanine nucleotide exchange factor (GEF) to the Rho GTPase and consequent effects on Mg^2+^-affinity of the latter (29). It is worth stating that our studies highlight a specific feature, the correlation of global closure with the alignment of catalytic elements to optimally engage Mg^2+^, that appears to be universal for all P-loop enzymes. However, additional processes that are diverse and dependent on context e.g., nucleotide selectivity (ATP vs GTP) (30), the presence of positively charge provided by an arginine finger (31) or a monovalent cation (32), exclusion of bulk solvent for a P-loop kinase (15) etc. are also necessary to form the corresponding reactive states.

The goal of the present analyses was to identify overarching structural features that characterize the active and inactive states of P-loop enzymes. As described above, we used a large library of crystal structures to identify specific structural features, local and global. We note that crystal structures, even in the numbers and the variety deployed here, in various liganded states, are imperfect proxies for the overall conformational landscape of P-loop enzymes. This is due to multiple factors e.g., use of cryogenic temperatures (33), crystal packing effects (34), and modeling errors (35), that introduce biases and/or “noise” into our datasets. Nevertheless, our present analyses provide guideposts to probe for specific features in future studies of P-loop enzymes ideally using a synergistic approach using experiment and computation as in the case of Wzc (14, 15).

## Materials and Methods

### Curation of structures

To construct a library of structures, the list generated by “ATPase” or “GTPase” keyword searches in the protein databank (PDB) were assembled and from these the ASCE ATPases, SIMIBI and TRAFAC GTPases were selected. The structural coordinates for individual entries were visually inspected with each relevant chain in the asymmetric unit being treated as a separate entity to account for conformational heterogeneity. NMR-determined structures were excluded and a resolution cutoff of 3.5 Å was used in the curation of structures. Assignment to the protein family type for each entry was determined from the corresponding annotation in the RCSB or from UniProt databases, and structures were classified accordingly. The bound nucleotide for each structure was also recorded with natural nucleotides and analogs (including transition state analogs) were treated as equivalent. Additionally, within each nucleotide-bound state, the Mg^2+^-bound and Mg^2+^-free states were separated. Other divalent cations (that were rarely present), such Ca^2+^ or Mn^2+^, were treated as equivalent to Mg^2+^. Additionally, active site mutants, and P-loop ATPases with significantly divergent Walker-A and Walker-B motifs (except Walker-B Asp/Glu variants), were excluded.

### Transformation of structures and projection onto the spherical coordinate frame

First, a given structure was translated to place the center-of-mass (COM1) of the C*α* atoms comprising the first turn of *α*^P^ (P1 in Fig. 2) at the origin. Next, a rotation was carried out to place the center-of-mass of the C*α* atoms of the second turn of *α*^P^ (COM2) on the z-axis (P2 in Fig. 2). Finally, a rotation about the z-axis was performed to place the C*α* atom of Ser_A_/Thr_A_ (P3 in Fig. 2) on the x-axis in the positive direction. The coordinates for the curated structures were utilized to compute the positions of the reference points P1, P2 and P3 (Fig. 2) that were subsequently utilized to obtain the corresponding *D*^W^, *ϕ* and *θ* values. *D*^W^ was obtained as the Euclidian distance between the Ser_A_/Thr_A_ C*β* and the Asp_B_ C*γ* (or C*δ* in case of a Glu_B_) atoms. The angle *ϕ* was the angle subtended by the vector connecting the origin to the C*α* atom of the Walker-B Asp 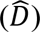 and a unit vector along the z-axis, 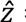 = (0, 0, 1):

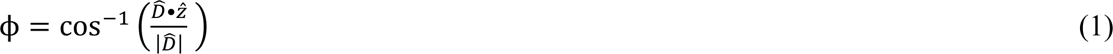

*θ* was defined as the angle between 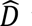 vector and a unit vector along the x-axis, 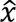 = (1, 0, 0):

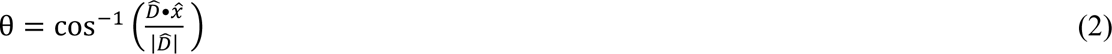

Additional distances *D^α^*, *D^β^*, *d^α^* and *d^β^* were obtained as Euclidian distances between atoms described in the text. All manipulations were carried out using in-house programs written in C++. For the projections of the REST2-generated conformational ensembles for the *E. coli* Wzc catalytic core (14) onto the *D*^W^*−ϕ−θ* coordinate frame, a modified version of the code that utilized the Pteros library (36) for handling GROMACS (37) trajectory files.

### Clustering and statistics

Mean Shift (MS) clustering (20, 38) in spherical coordinates was performed in the R programing language using the local principal curve methods (LPCM) library and analyzed utilizing built-in functionalities within R. The procedure produced several clusters where the majority contained only few structures. A cluster was accepted for analysis if it consisted of more than 9 structures for the KG group or 10 structures for the ASCE group, where the structures originate from at least 2 different proteins including homologs of a selected protein from a different organism. Following the clustering procedure, the individual clusters were visually inspected and assigned as open or closed based on the cluster centers for *D*^W^.

## Acknowledgements

This work was supported by NSF grant MCB1937937. The authors thank Dr. Sébastien Alphonse (CCNY) for his critical comments on this manuscript.

## Data availability

Details of the structural datasets and characteristics including PDB files used in the analysis and corresponding annotations are provided in Datasets S1 and S2. All other data used in the study are included in the article an *SI Appendix*.

## Author contributions

F. H. and R. G. conceived the project; F. H. developed the methodology and performed all the analyses. F. H. prepared a first draft of the paper and figures that were refined by R. G. with input from F. H.

## Competing interests

The authors declare that they have no competing interests.

## Supplementary Material for

**Fig. S1.**
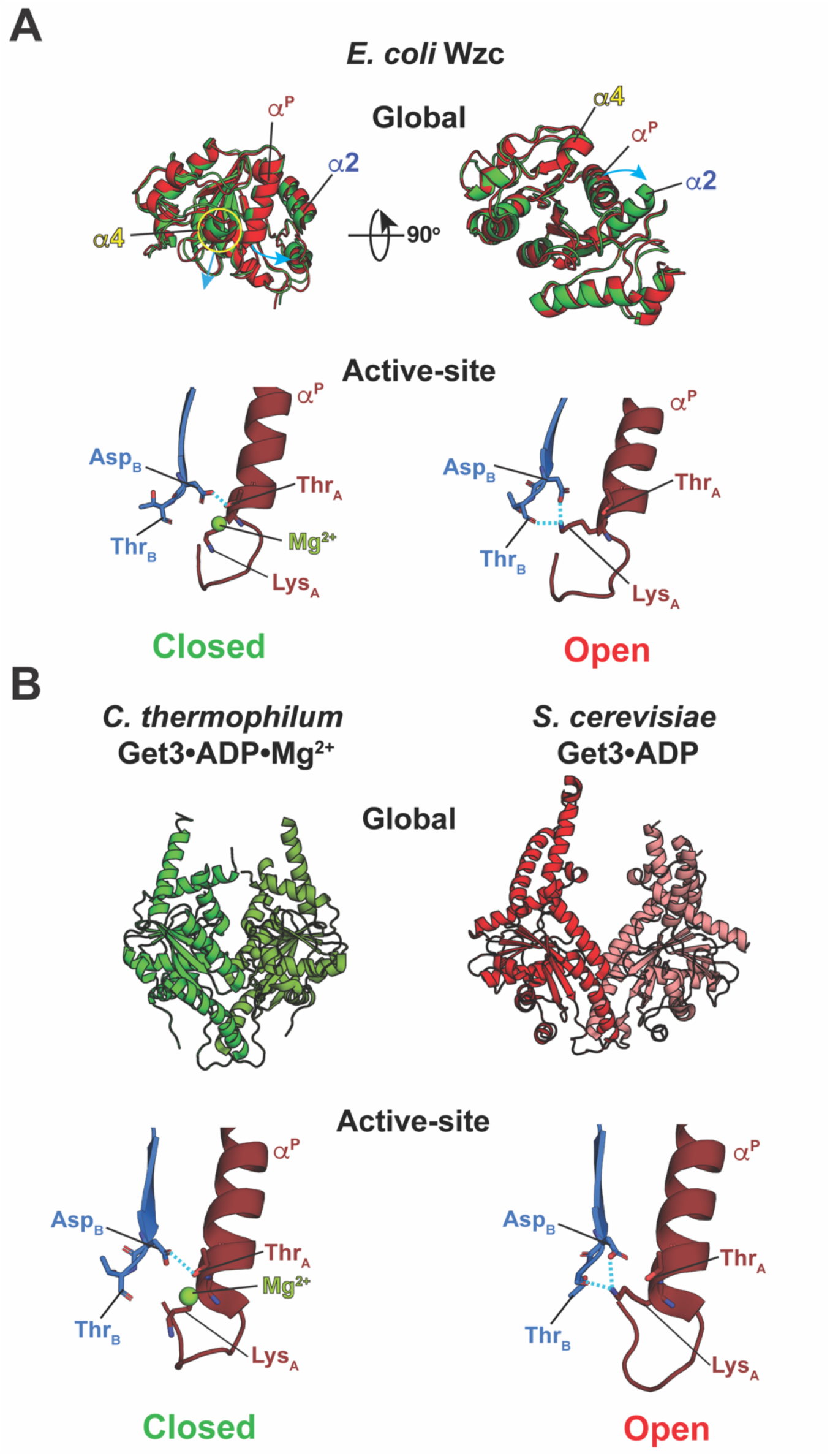
Characteristics of global open and closed states and their correlation with active-site conformation in P-loop enzymes. **(A)** Global transitions (top panel) and the corresponding changes in interactions at the active site (lower panel) are shown using representative structures drawn from the open (red) and closed (green) state ensembles of the catalytic core of *E. coli* Wzc in its complex with ATP•Mg^2+^ (1). Global opening occurs by a downward displacement of helix *α*4 (labeled in yellow), and an outwards rotation of the *α*^P^ (labeled in brown) and *α*2 (labeled in blue) helices with respect to the protein core (cyan arrows). These global changes are coupled with a reshuffling of active-site interactions (lower panel) where the hydrogen-bond between the sidechains of Asp_B_ and Thr_A_, required to coordinate Mg^2+^, is lost in the open state, and the former now interacts with the Lys_A_ sidechain. The Lys_A_ sidechain also interacts with the backbone of a residue (Thr_B_) that immediately follows Asp_B_ in the open state. **(B)** The structure of *C. thermophilum* Get3 forms a closed state in its complex with ADP•Mg^2+^ (PDB: 3IQX, green) (2); the absence of Mg^2+^ leads to an open state represented by the structure of *S. cerevisiae* Get3 in its complex with ADP (PDB:3A37, red) (3). The reshuffling of the active-site interactions seen in Get3 is identical to that seen for Wzc. The bound nucleotides are not shown in any of the cases.

**Fig. S2.**
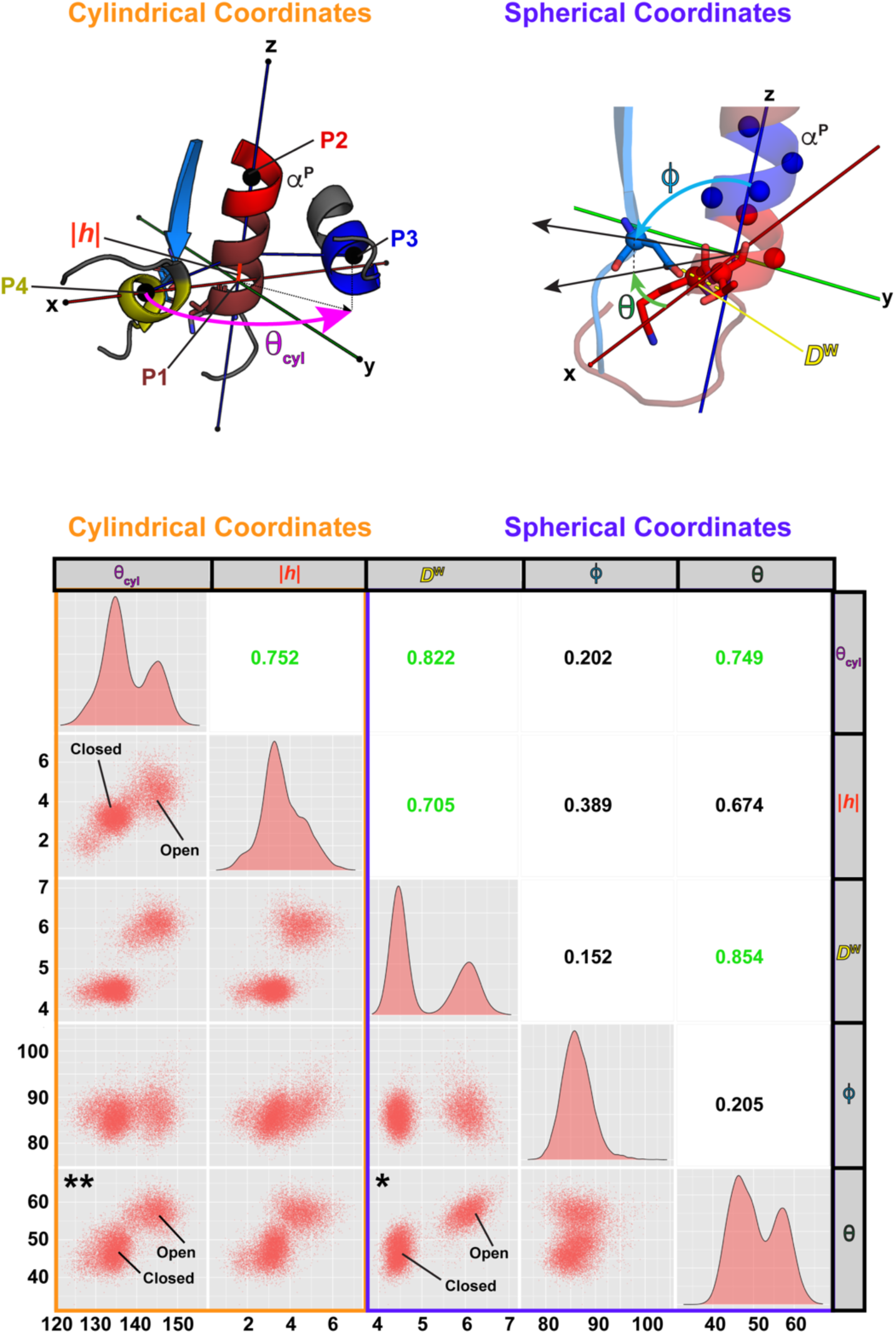
Validation of the spherical coordinate frame. The cylindrical coordinate frame (top left panel) involving a rise (|*h*|) and an angle *θ*_cyl_ can distinguish between global open and closed states in the structural ensemble of the catalytic core of the BY-kinase Wzc in its complex with ATP•Mg^2+^ (1). This cylindrical coordinate frame relies on an alignment of the *α*^P^ helix about the z-axis via points P1 (the center of mass of the C*α* atoms of the residues comprising the first two turns of *α*^P^), and P2 (center of mass of the C*α* atoms of the residues comprising the third turn of *α*^P^). Additionally, the BY-kinase-specific structural elements, the helices *α*2 (for P3) and *α*4 (for P4), define an angle *θ*_cyl_ measured as the counterclockwise rotation of P3 with respect to the protein core (magenta arrow). The rise |*h*| is measured as the absolute displacement (red bar) of point P4 with respect to the core. The current spherical coordinate system (also see Fig. 2) is also shown on the top right panel for reference. The *θ*_cyl_ and |*h*| distributions are multimodal serving to resolve the open and closed states as shown before (1, 4). Projection of these same results onto the newly defined spherical coordinate frame also resolves the open and closed states. A 5-by-5 correlation matrix between pairs of variables (*θ*_cyl_, *|h|*, *D*^W^*, θ* and *ϕ*) is shown on the bottom panel. The diagonal elements show the one-dimensional distributions for a particular variable. The upper triangular matrix elements show the pairwise Pearson correlation coefficients (values greater than 0.7, indicating high degree of correlation, are labeled in green); the lower triangular elements illustrate pairwise point projections of the ensemble. The spherical coordinate frame reliably captures the open and closed states (see for example the *D*^W^ vs *θ* plot indicated by the “*”) and for Wzc achieves a clearer separation between them compared to the cylindrical coordinate frame. Note that *in the specific case* of Wzc, the distribution of *ϕ* is unimodal and shows poor correlation with *D*^W^. Inspection of the cross terms between spherical and cylindrical coordinates reveals excellent correlation between states in the two reference frames (e.g., *θ*_cyl_ vs *θ*, indicated by the “**”, with a correlation coefficient of 0.749). This comparison shows that the spherical coordinate frame not only quantifies rearrangements local to the active site but also provide an excellent measure of the global conformational changes without the need for a class/family-specific definitions.

**Fig. S3.**
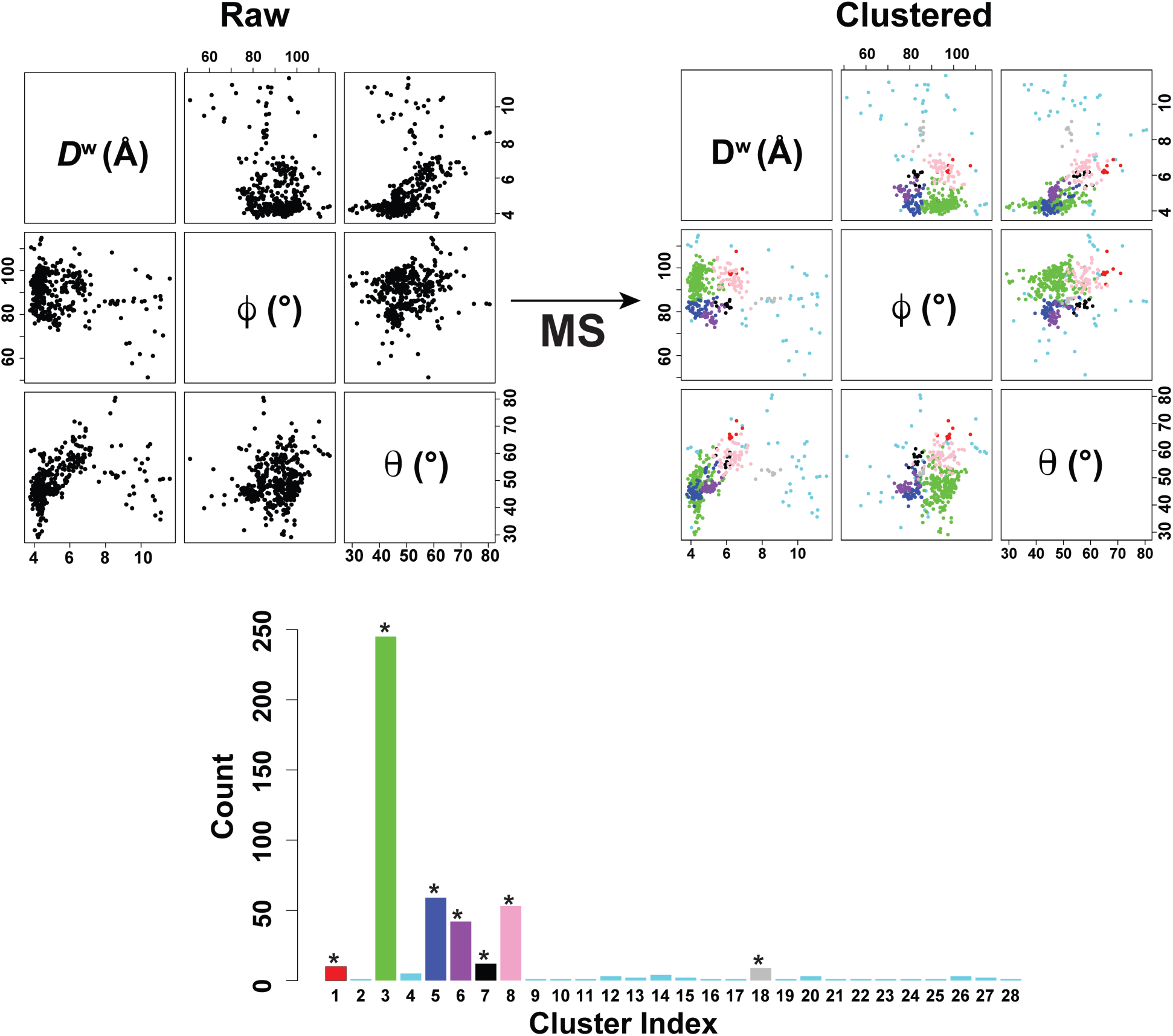
Clustering of the KG structures projected onto the spherical coordinate frame using the Mean Shift algorithm. The upper left panel shows pairwise point projections of the data set across the three coordinates -*D*^W^, *ϕ*, and *θ*. The same plot is shown on the upper right panel after clustering using the Mean Shift (MS) algorithm (5, 6). The MS clustering procedure relies on only a single free parameter known as the bandwidth, which in this case, was chosen as 5% of the data range for each variable. The lower panel shows the number of structures in each cluster color coded using the same color scheme as in the upper right panel and throughout the text. The clusters indicated by the ‘*’ are discussed in the text. Clusters with low structure counts (< 9), colored cyan, span a wide and diffuse range without any specific centering (upper right panel). These latter clusters are not discussed further since their low structure count does not allow for the extraction of meaningful statistics. It should be noted, however, that the diffuse nature of the cyan points is occasionally indicative of a partially unfolded Walker-B motif.

**Fig. S4.**
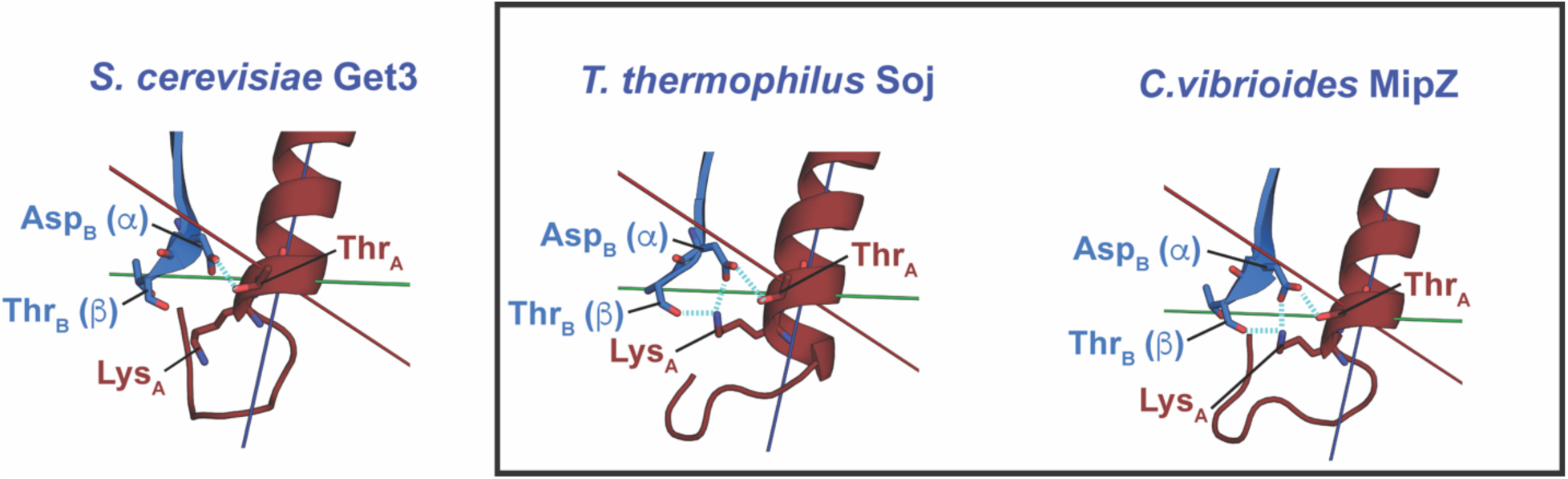
Intermediate states in the dark-blue cluster of KG structures. Closed state structures normally display the characteristic interaction between the sidechains of Asp_B_ and Thr_A_; the Lys_A_ sidechain is in a downward orientation and disengaged from both the Asp_B_ sidechain and the backbone of the residue at the *β*-position leading to large values for *D^α^* (∼7 Å) and *D^β^* (∼4.5 Å) as in *S. cerevisiae* Get3 (left panel; PDB: 2WOJ, chain A; the bound Mg^2+^ ion is not shown) (7). In contrast, anomalous structures of the dark-blue cluster represent intermediate configurations in which the LysA sidechain interacts with Asp_B_ and the *β*-position leading to short *D^α^* (∼4 Å) and *D^β^* (∼2.5 Å) distances while simultaneously displaying the Asp_B_/Thr_A_ interaction. These structures are Mg^2+^ free, but Lys_A_ has not acquired a downward orientation as expected for an open state structure. Examples of these intermediate structures (shown within the rectangle) include *T. thermophilus* Soj (middle panel; PDB: 1WCV, chain A) (8) and *C. vibrioides* MipZ (right panel; PDB: 2XJ4, chain A) (9).

**Fig. S5.**
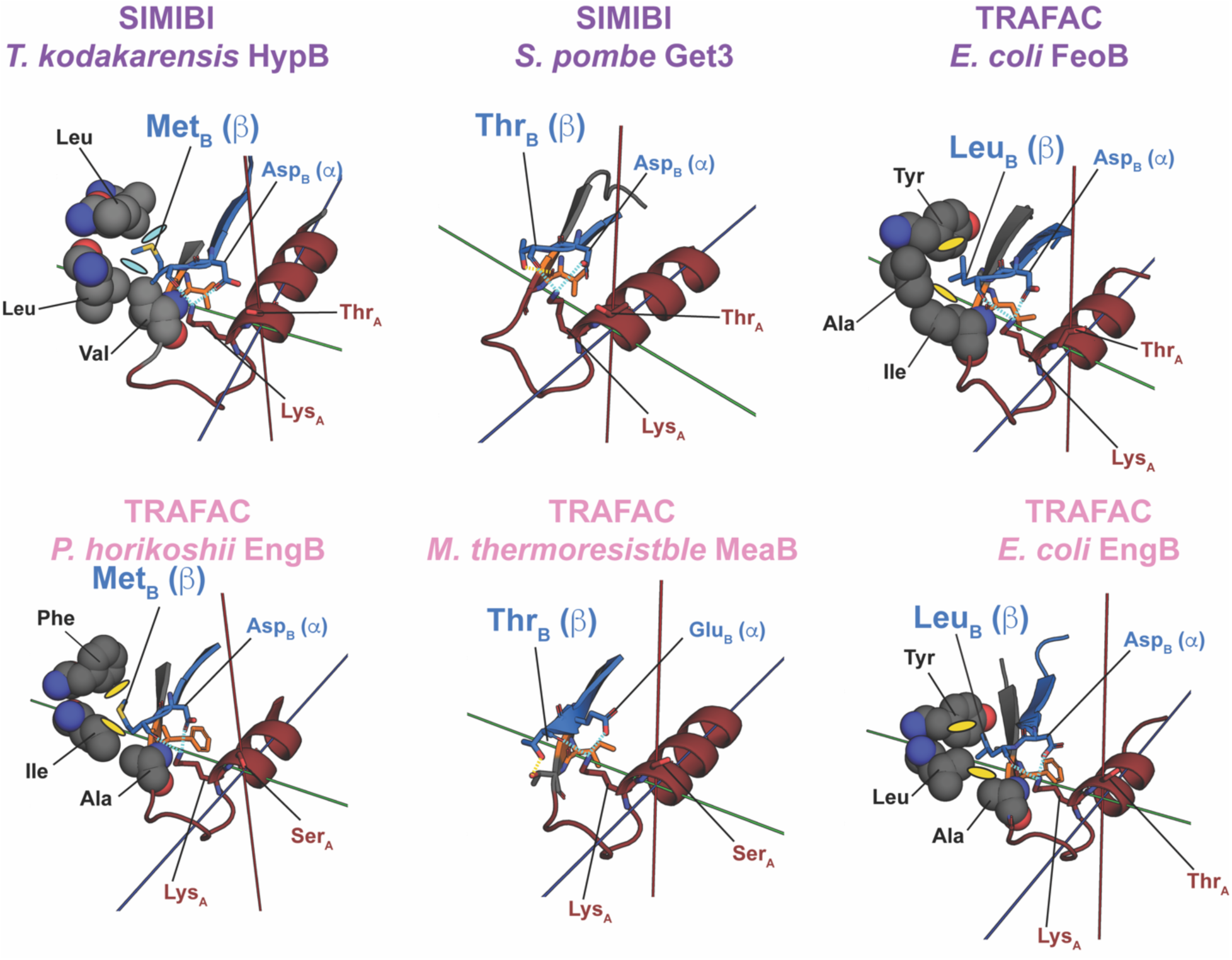
The role of the residue at the *β*-position in stabilizing the open state. Representative structures illustrating the interaction profile between Lys_A_ and the residue immediately following Asp_B_ i.e., the residue at the *β−*position for structures drawn from the purple and pink clusters in Fig. 3. Examples depict interactions involving the most commonly appearing residues at this position, namely Thr, Met and Leu (see Fig. 4C). In all cases, the *β*-residue is stabilized through range of interactions including hydrogen-bonds (in case of Thr; yellow dashes) or hydrophobic interactions (e.g., Met, Leu; yellow ovals) with spatially proximal residues outside the Walker motifs (remote hydrophobic residues shown in space filling representation). Examples shown include *T. kodakarenis* HypB (PDB: 5AUQ, chain F) (10), *S. pombe* Get3 (PDB: 2WOO, chain A) (7), *E. coli* FeoB (PDB: 3I8S, chain C), *P. horicoshii* EngB (PDB: 2CXX, Chain A), *M. thermoresistible* MeaB (PDB: 3TK1, chain A) (11), and *E. coli* EngB (PDB: 1PUI, chain A).

**Fig. S6.**
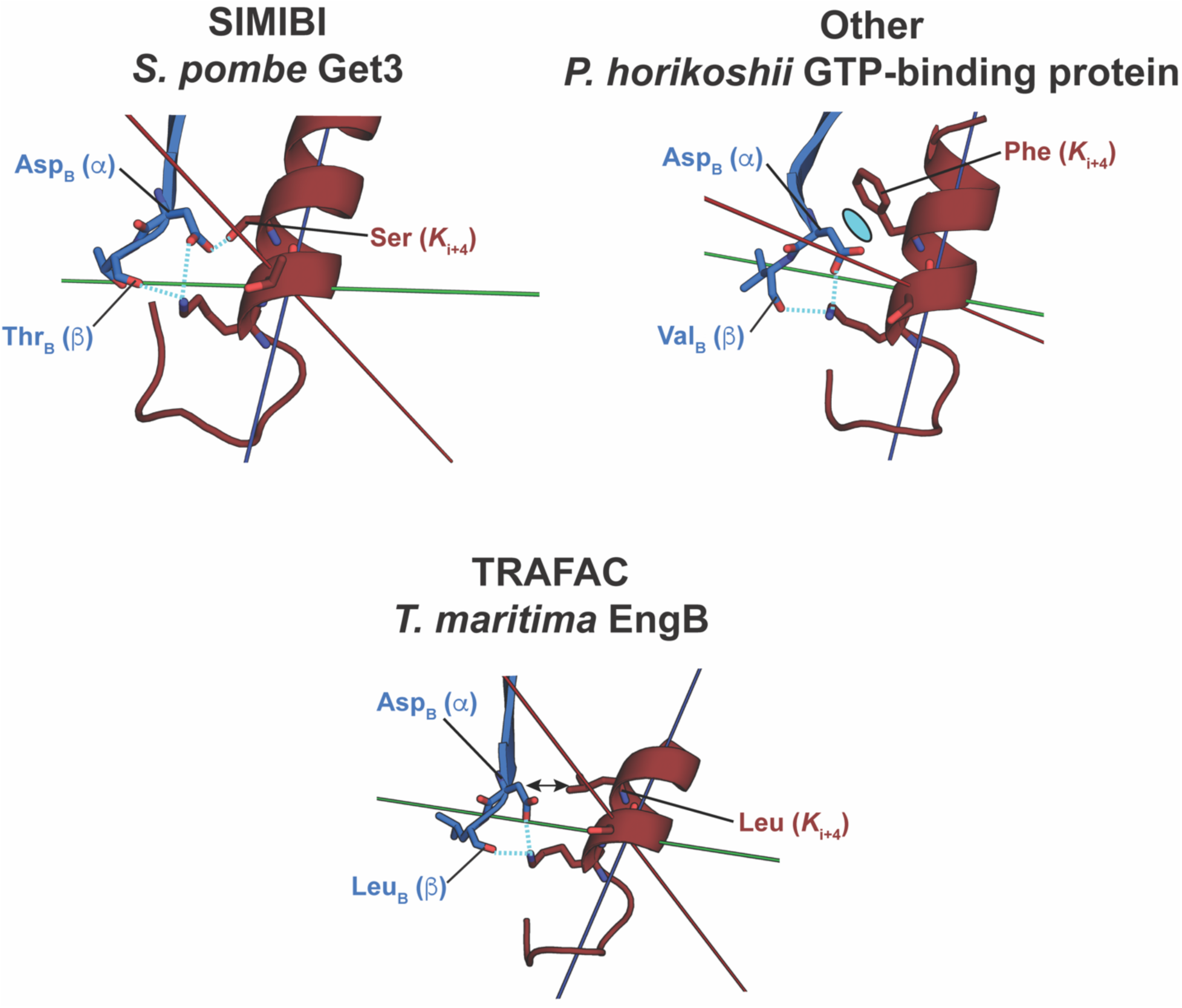
Role of a key *α*^P^ residue in stabilizing the open state in KG structures. Specific examples of the interaction of the *α*^P^ residue at the 4^th^ position from Lys_A_ (*K*_i+4_) in stabilizing the open state through an interaction with Asp_B_ are shown on the top panel. These include a hydrogen-bond (cyan dashed lines) with a Ser in *S. pombe* Get3 (PDB: 2WOO, chain A) (7) or anion-*π* interactions (cyan oval) with the *P. horikoshii* GTP binding protein (PDB: 1WXQ, chain A). In the presence of such interactions closed to open transitions involve changes in *ϕ*. In the absence of such interactions e.g., in *T. maritima* EngB (lower panel; PDB: 3PQC, chain A) (12), where this position is a Leu, opening can involve changes in *θ*.

**Fig. S7.**
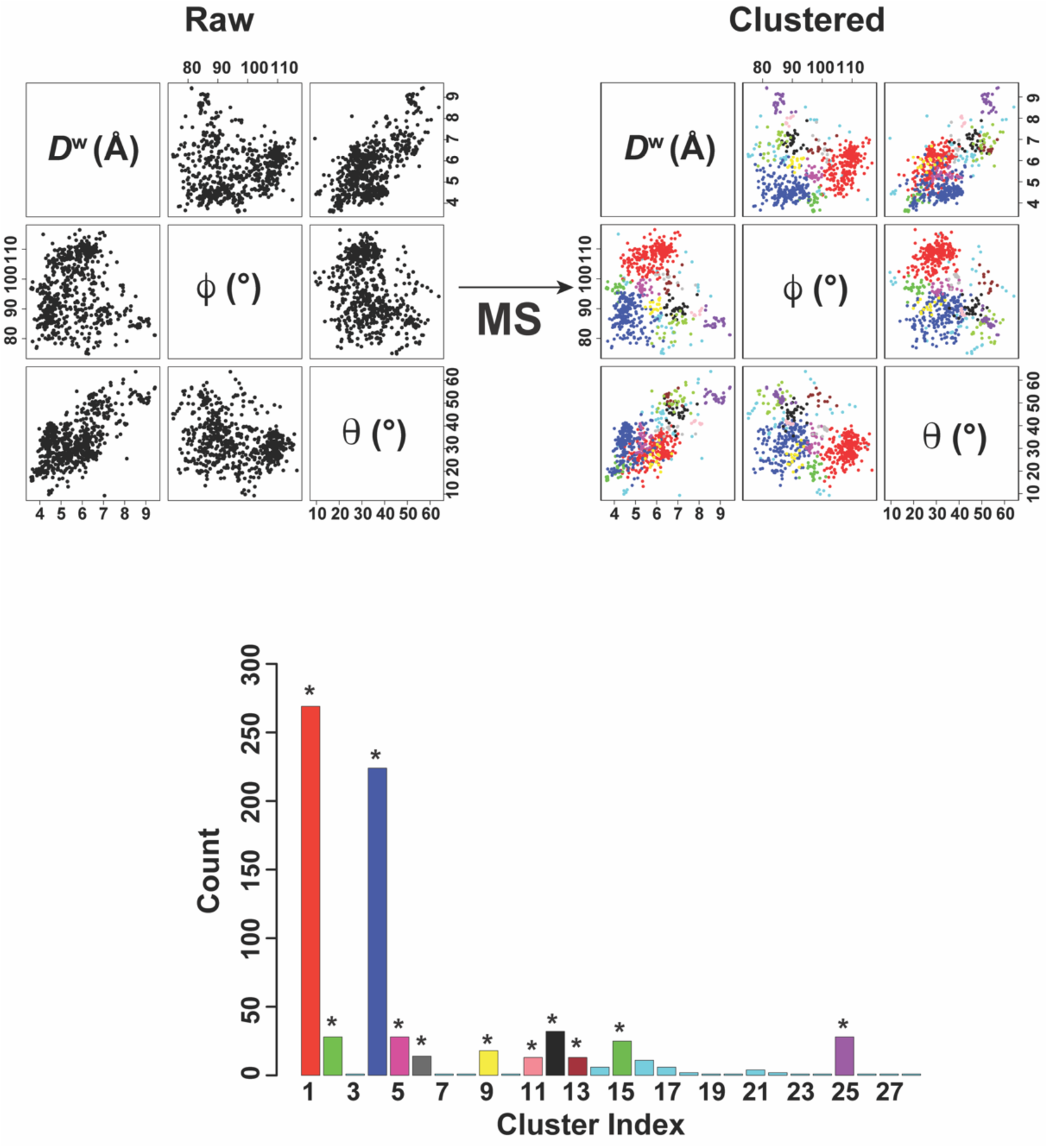
MS clustering of the ASCE structures in the spherical coordinate frame. The upper left panel shows pairwise point projections of the ASCE structure data set across the three coordinates *D*^W^, *ϕ*, and *θ*, and the right panel shows the same after MS clustering (5, 6). A bandwidth of 5% of the data range was used for each variable. The lower panel shows the structure count for each cluster color coded using the same color scheme as in the upper right panel and throughout the text. The clusters indicated by the ‘*’ are discussed in the text. Clusters with low structure counts (<10), colored cyan are not discussed further.

**Fig. S8.**
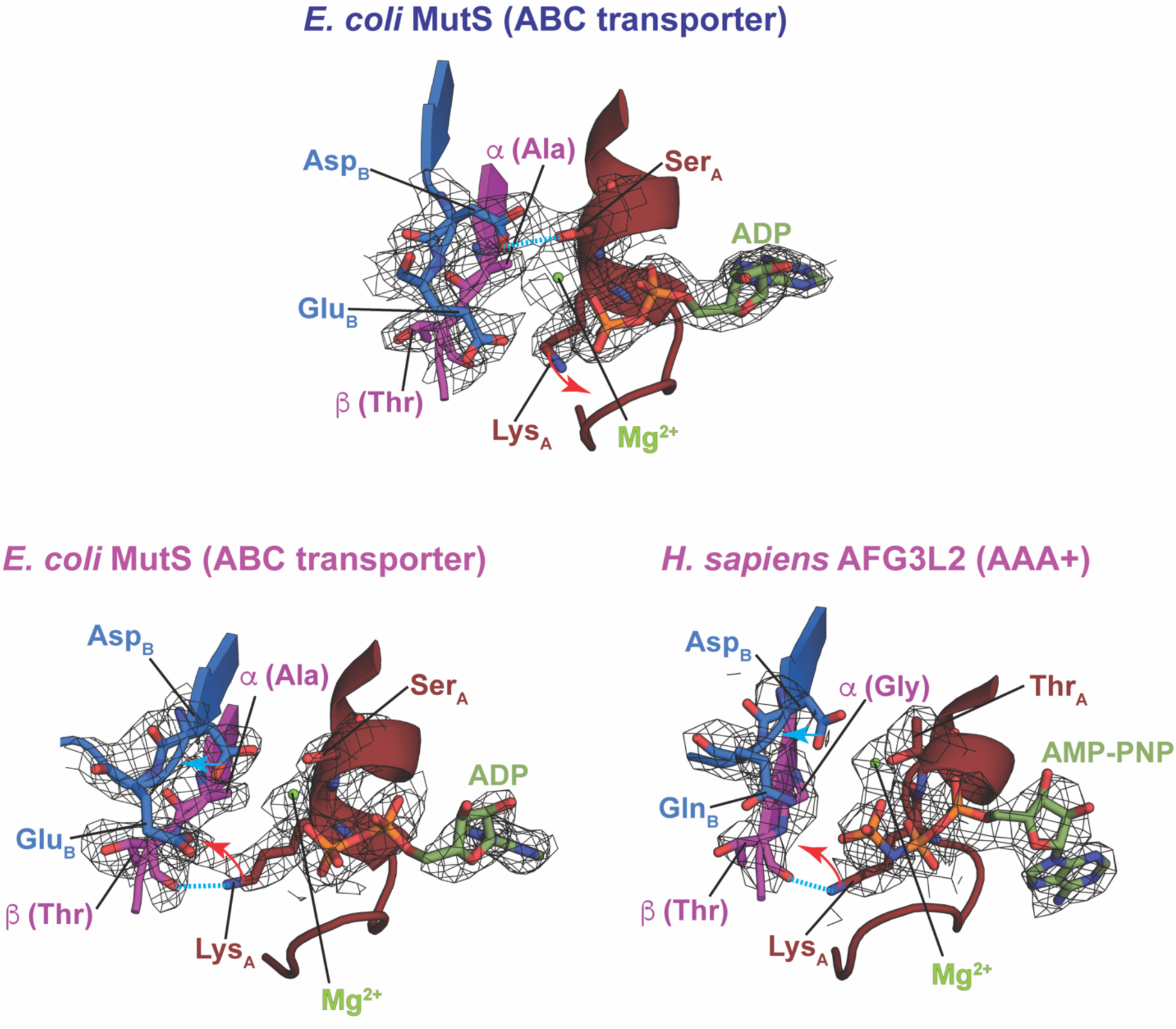
Conformation of key elements in the Mg^2+^-bound ASCE structures from the magenta cluster. **(A)** As shown by the structure of *E. coli* MutS (top panel; PDB: 1NG9, chain A) (13), in the closed state (dark-blue cluster in Fig. 6) Lys_A_ is displaced away (red arrow), the Asp_B_-Ser_A_ hydrogen-bond is established (cyan dashed line) and Mg^2+^ is bound. As shown for two Mg^2+^-bound structures of the open state drawn from the magenta cluster in Fig. 6, *E. coli* MutS (bottom left panel; PDB: 1NG9, chain B) and *H. sapiens* AFG3L2 (bottom right panel; PDB: 6NYY, chain D) (14), while the characteristic hydrogen-bond between the Lys_A_ sidechain and the carbonyl oxygen of a residue (*β*-position) on the additional strand (*β*4) is established (cyan dashed line) and Asp_B_ has moved away (cyan arrow) thereby breaking its contact with Ser_A_/Thr_A_, the Mg^2+^ ion has not yet been ejected. These likely represent intermediate states prior to complete opening and loss of Mg^2+^.

**Fig. S9.**
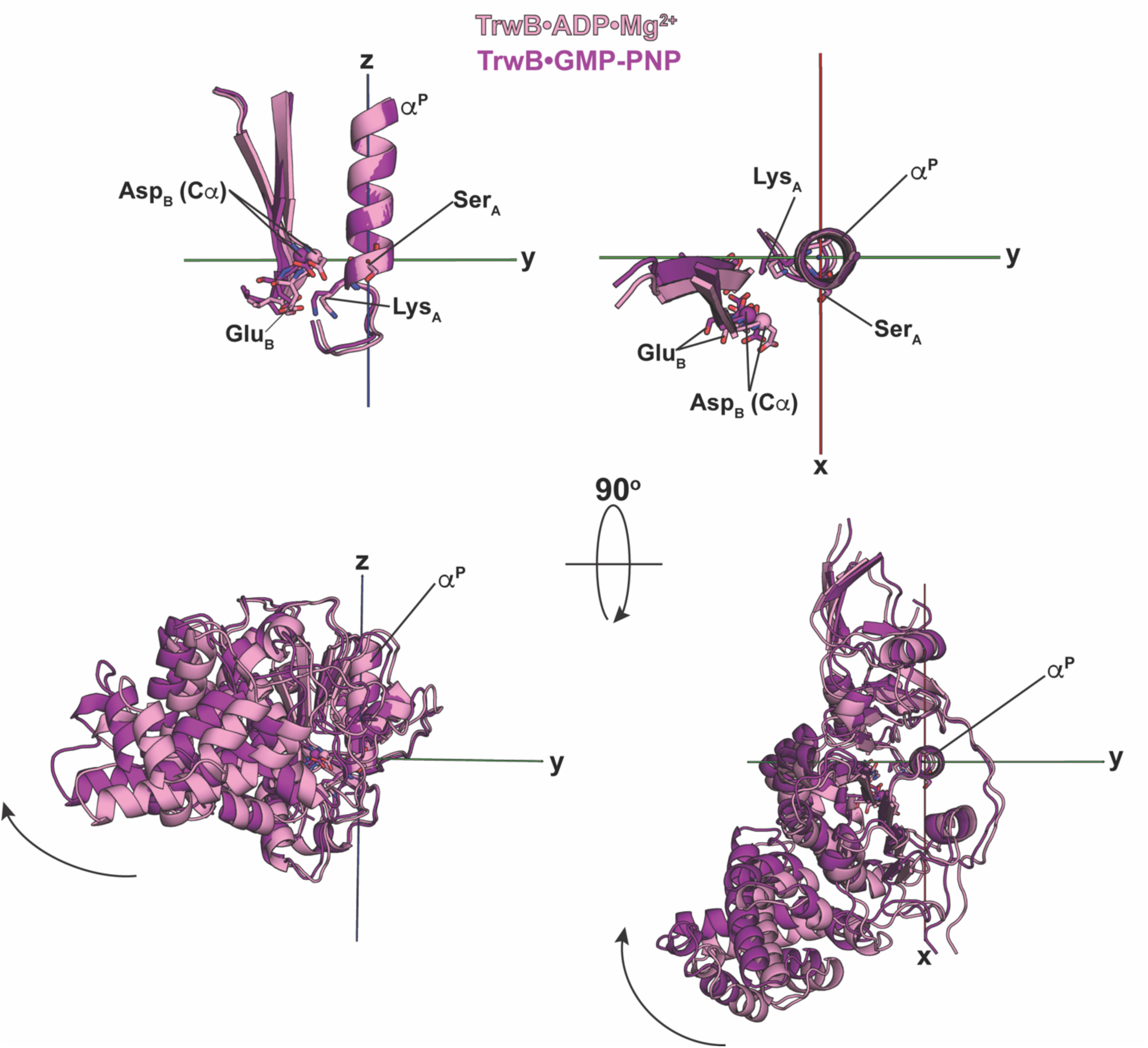
Conformation of key elements in a Mg^2+^-bound ASCE structure in the pink cluster. Shown are two structures of *E. coli* TrwB bound to ADP•Mg^2+^ (pink cluster in Fig. 6; PDB: 1GKI, chain A) (15) or to GMP-PNP (purple cluster in Fig. 6; PDB: 1GL6, chain A). While both are classified as open based on large *D*^W^ values, the former displays many characteristics of a closed state.

**Fig. S10.**
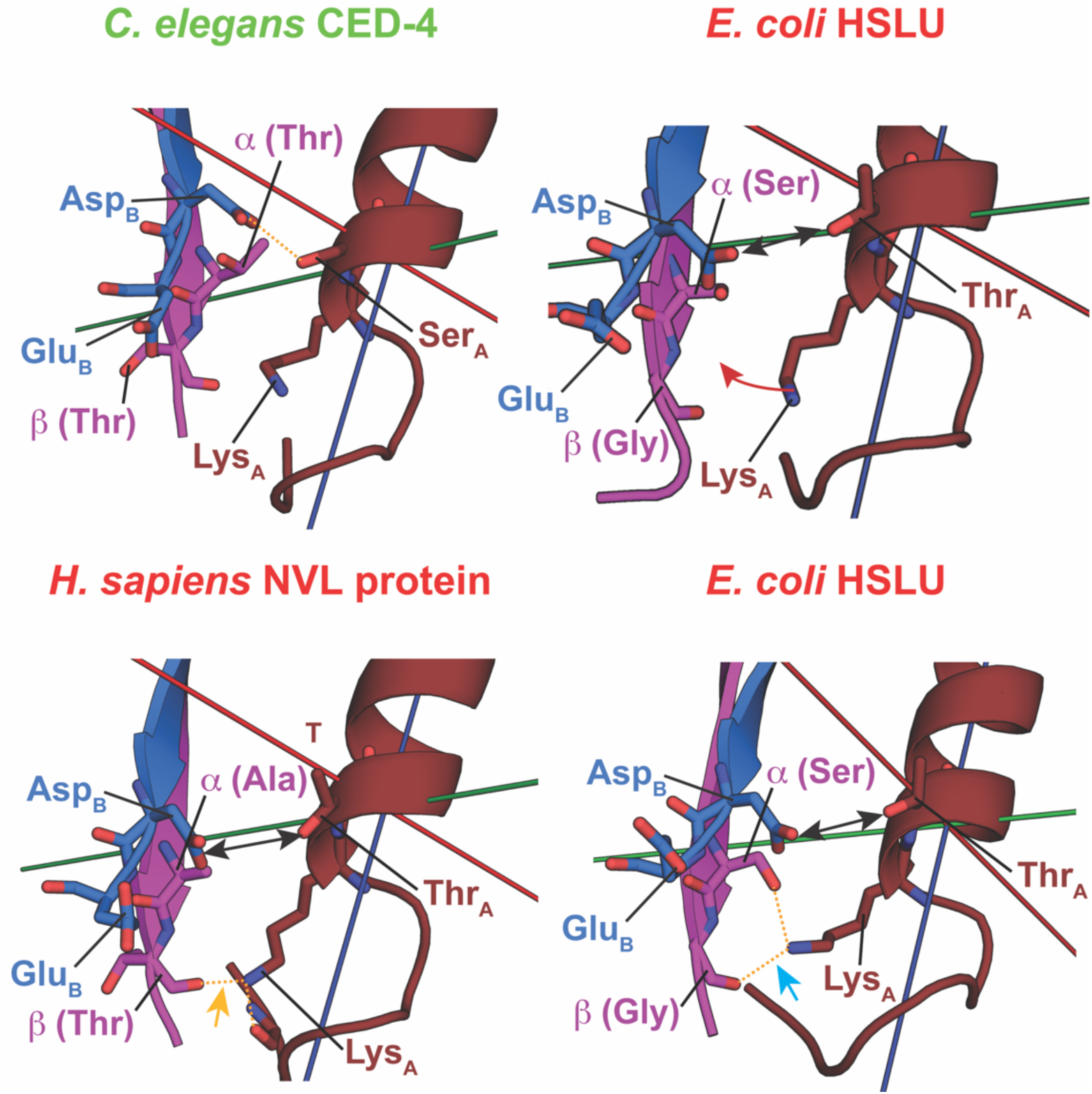
Role of the *α−* and *β−*positions in stabilizing the open states of AAA+ ATPases. In a closed state structure drawn from the green cluster in Fig. 6 (*C. elegans* CED-4, PDB: 2A5Y, chain A) (16), the Lys_A_ sidechain points away from the *α−* (Thr) and *β−*positions (Thr) positions. This contrasts a structure drawn from the red cluster (open state) in Fig. 6 (*E. coli* HslU, PDB: 1DO0, chain F) (17) in which the Lys_A_ sidechain is partially displaced (indicated by the red arrow; also in the distribution in Fig. 8B) towards the *α−* (Thr) and *β−*position (Gly) equivalents. However, the relevant distances are not optimal for the formation of characteristic hydrogen bonds as described for the KG class (see Fig. 4A). Hydrogen-bond formation between the Lys_A_ sidechain and the *β* (Thr) backbone oxygen (*H. sapiens* NVL protein, PDB: 2X8A, chain A; orange arrow here and in Fig. 8C) or with both the *α−*sidechain and the *β−*backbone (*α*: Ser, *β*: Gly; *E. coli* HslU, PDB: 1HT1, chain I; cyan arrow here and in Fig. 8B) (18) is occasionally seen.

**Fig. S11.**
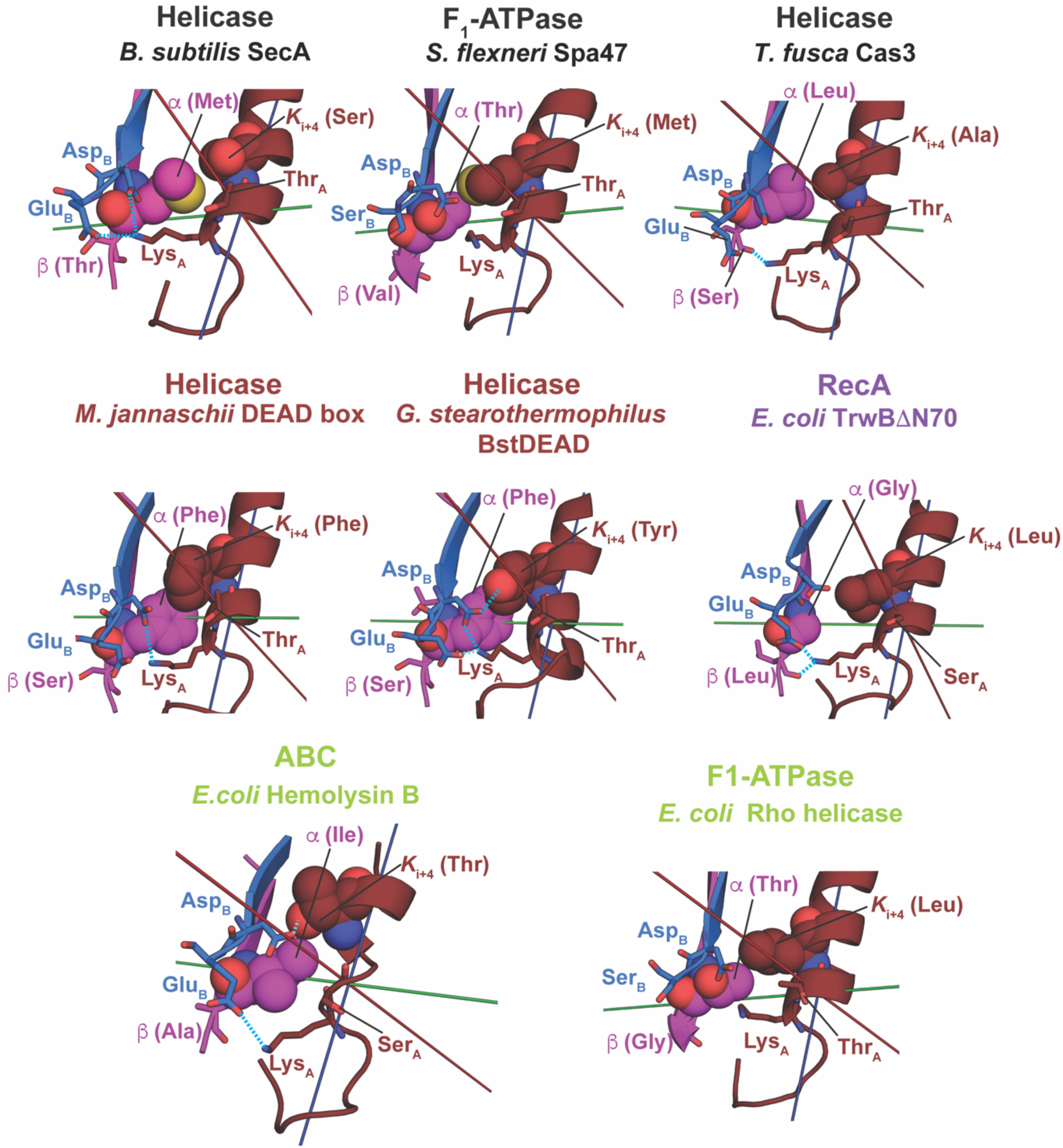
Importance of the *K*_i+4_ residue in maintaining the open state in non-AAA+ ASCE ATPases. Several examples of open state structures from the helicase, F_1_-ATPase, RecA and ABC transporter families are shown. The structures shown include *B. subtilis* SecA (PDB: 1TF5, chain A) (19), *S. flexneri* Spa47 (PDB: 5SWJ, chain A) (20) and *T. fusca* Cas3 (PDB: 4QQW, chain A) (21) all from the black cluster in Fig. 6; *M. jannaschii* DEAD box protein (PDB: 1HV8, chain A) (22) and *G. stearothermophilus* BstDEAD (PDB: 1Q0U, chain A) (23) from the brown cluster in Fig. 6; *E. coli* TrwB (PDB: 1E9S, chain A) (15)from the purple cluster in Fig. 6; *E. coli* hemolysin B (PDB: 1MT0, chain A) (24) and *E. coli* Rho (PDB: 2HT1, chain A) (25)from the lime cluster in Fig. 6. It is evident that a direct interaction between the a-and *K*i+4 residues is possible in all cases except in RecA that contains a glycine at the former position.

**Fig. S12.**
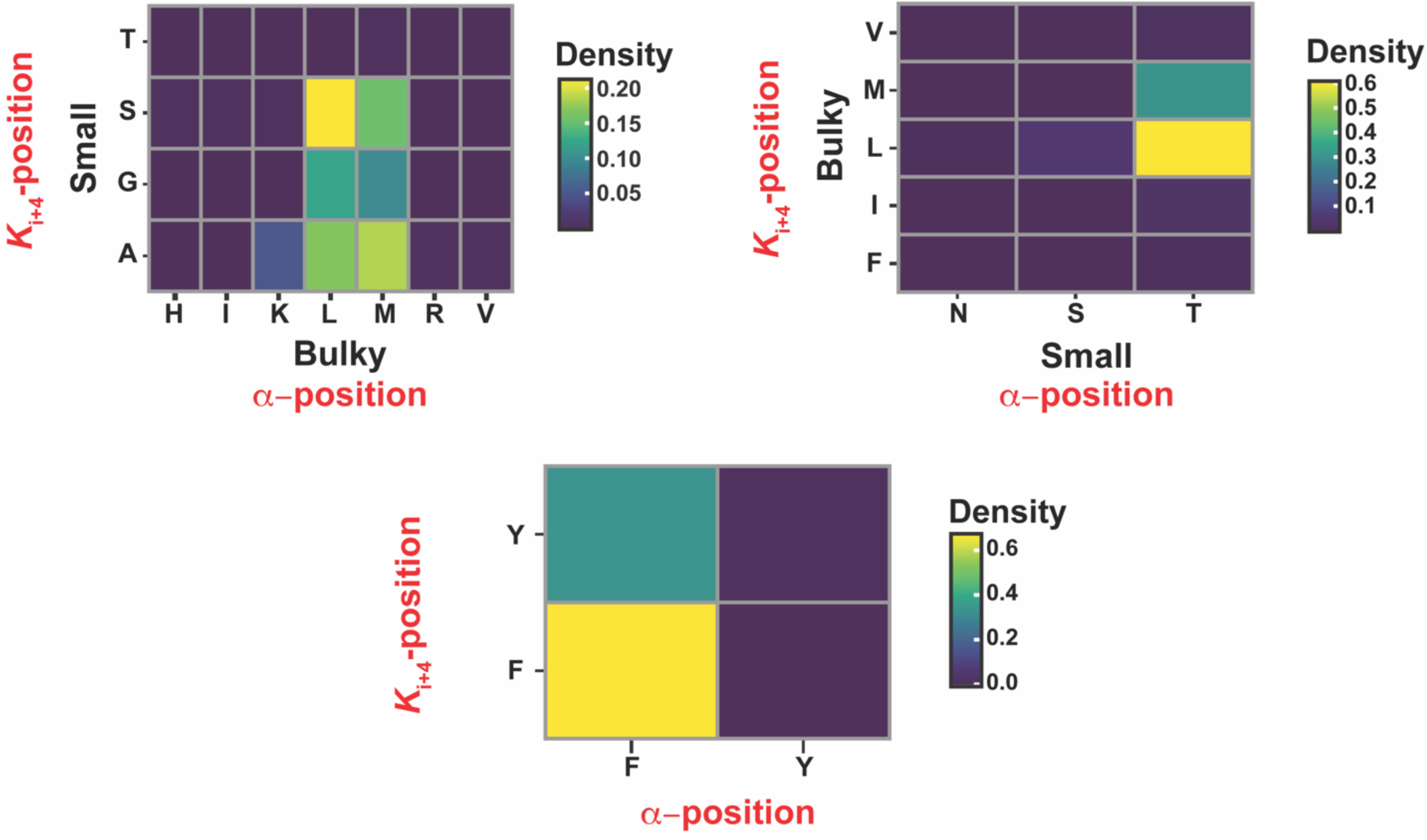
Pairwise correlation of amino acid types in the *α* and *K*_i+4_ positions. The correlation plots were constructed from 500 representative sequences through individual BLAST searches (75% identity cutoff) using *B. subtilis* SecA (top left panel), or *S. flexneri* Spa47 (top right panel), or *G. stearothermophilus* BstDEAD (lower panel) sequences. The amino acid identity of the *α−* and *K*_i+4_ positions were extracted from aligned sequences and pairwise occurrence of amino acid types were plotted as density plots. In each case the x- and y-axis represent the range of observed amino acid types for each alignment. Note in the case of SecA (top left) the search sequence contained an M/S for the *α*/*K*_i+4_ pair while the BLAST produced a comparable most likely pair of L/S, and in general the variance preserves a bulky/small pair for the *α* and *K*_i+4_ positions respectively. In the case of *S. flexneri* Spa47 (top right) the search sequence contained a T/M pair and in that case again the most common occurrence was a comparable T/L pair, i.e., a small/bulky amino acid for the respective positions. In the case of *G. stearothermophilus* BstDEAD (lower panel) an F/Y pair in the search sequence, produced a most commonly F/F pair. This analysis reflects the overall trends (shown on the bottom panel of Fig. 9A) determined by tallying the *α−* and *K*_i+4_ positions for the ASCE structures used in the analysis.

**Table S1.** Characteristics of the KG clusters.

| Cluster | $D^w$ center (Å) | $\phi$ center (°) | $\theta$ center (°) | $D^w$ SD (Å) | $\phi$ SD (°) | $\theta$ SD (°) |
| --- | --- | --- | --- | --- | --- | --- |
| Red | 6.3 | 97 | 63 | 0.2 | 4 | 2 |
| *Green* | 4.3 | 97 | 47 | 0.4 | 5 | 5 |
| *Dark-blue* | 4.2 | 84 | 46 | 0.4 | 2 | 4 |
| Purple | 4.9 | 78 | 46 | 0.3 | 3 | 2 |
| Black | 6.0 | 83 | 57 | 0.3 | 2 | 2 |
| Pink | 6.5 | 93 | 58 | 0.5 | 5 | 3 |
| Gray | 8.6 | 85 | 52 | 0.4 | 1 | 1 |

**Table S2.** Characteristics of the ASCE clusters.

| Cluster | $D^w$ center (Å) | $\phi$ center (°) | $\theta$ center (°) | $D^w$ SD (Å) | $\phi$ SD (°) | $\theta$ SD (°) |
| --- | --- | --- | --- | --- | --- | --- |
| Red | 6.1 | 109 | 31 | 0.6 | 3 | 5 |
| *Green* | 4.2 | 97 | 22 | 0.3 | 2 | 3 |
| *Dark-blue* | 4.5 | 90 | 37 | 0.5 | 4 | 6 |
| Magenta | 5.2 | 96 | 32 | 0.2 | 2 | 4 |
| Gray | 6.5 | 101 | 39 | 0.5 | 2 | 1 |
| Yellow | 5.9 | 90 | 26 | 0.3 | 2 | 4 |
| Pink | 7.8 | 88 | 42 | 0.2 | 1 | 1 |
| Black | 7.1 | 90 | 55 | 0.4 | 2 | 4 |
| Brown | 6.5 | 97 | 53 | 0.2 | 3 | 2 |
| Lime | 6.9 | 87 | 49 | 0.5 | 3 | 4 |
| Purple | 8.8 | 85 | 52 | 0.3 | 2 | 3 |

Clusters classified as closed are shown in *italics*.
Center: The center represents the most probable value for a given variable in each cluster.
SD: Standard deviation for a given variable in each cluster.

## Notes

### Competing Interest Statement

The authors have declared no competing interest.

